# Unusual lineage plasticity revealed by YY1 knockout in pro-B cells

**DOI:** 10.1101/2024.03.22.586298

**Authors:** Sarmistha Banerjee, Sulagna Sanyal, Suchita Hodawadekar, Sarah Naiyer, Nasreen Bano, Anupam Banerjee, Joshua Rhoades, Dawei Dong, David Allman, Michael L. Atchison

**Affiliations:** Department of Biomedical Sciences, School of Veterinary Medicine, University of Pennsylvania, 3800 Spruce Street, Philadelphia, PA 19104; Department of Pathology and Laboratory Medicine, Perelman School of Medicine, University of Pennsylvania, 3400 Civic Center Blvd, Philadelphia, PA 19104; Merck & Co, Inc., 351 North Sumneytown Pike, North Wales, PA 19454

**Keywords:** YY1, B cell, lineage commitment, lineage plasticity, sc-RNA-seq, alternative lineages, transcription

## Abstract

During B cell development, cells progress through multiple developmental stages with the pro-B cell stage defining commitment to the B cell lineage. YY1 is a ubiquitous transcription factor that is capable of both activation and repression functions. We find here that knockout of YY1 at the pro-B cell stage eliminates B lineage commitment. YY1 knockout pro-B cells can generate T lineage cells *in vitro* using the OP9- DL4 feeder system, as well as *in vivo* after injection into sub-lethally irradiated Rag1^-/-^ mice. These T lineage-like cells lose their B lineage transcript profile and gain a T cell lineage profile. Single cell-RNA-seq experiments showed that as YY1 knockout pro-B cells transition into T lineage cells, various cell clusters adopt transcript profiles representing a multiplicity of hematopoietic lineages indicating unusual lineage plasticity. Given the ubiquitous nature of YY1 and its dual activation and repression functions, YY1 likely regulates commitment in multiple cell lineages.

## Introduction

Development of the B lymphocyte lineage follows an ordered progression of cell stages including common lymphoid progenitors (CLPs), pre-pro-B cells, pro-B cells, pre-B cells, immature B cells, and additional more mature and specialized B cells (Henderson and Calame 1998; Bartholdy and Matthias 2004; Shapiro-Shelef and Calame 2005; Hagman and Lukin 2006; Pang et al. 2014; Miyai et al. 2018). Precursor cells (pre-pro-B and earlier) are capable of adopting alternative lineage fates, but lineage commitment occurs at the pro-B cell stage subsequent to somatic rearrangement of immunoglobulin heavy chain (IgH) gene variable, diversity, and joining (VDJ) segments to produce a functional IgH gene (Nutt et al. 1999; Jung et al. 2006).

During the transition to committed pro-B cells, specific transcription factors function to initiate gene expression profiles necessary for development of B lineage cells, while simultaneously repressing the expression of genes needed for the development of alternative lineages (Bain et al. 1994; Nutt et al. 1997; Henderson and Calame 1998; Su et al. 2003; Busslinger 2004; Hagman and Lukin 2006; Liu et al. 2007; Banerjee et al. 2016; Kleiman et al. 2016). Transcription factors EBF1 and E2A are crucial regulatory proteins that initiate early stages of B lineage development, enabling expression of key genes such as Igll1, VpreB1, Cd79a, Rag1, and Rag2, as well as expression of transcription factor Pax5. Pax5 further strengthens the B lineage pathway by i) stabilizing the B lineage expression profile, ii) repressing the expression of alternative lineage genes, iii) inducing IgH V-to-DJ rearrangements and, iv) initiating locus contraction of the IgH locus (Kee and Murre 2001; Fuxa et al. 2004). Upon executing these regulatory networks at the pro-B cell stage, the cells are exclusively committed to the B cell lineage and cannot adopt other lineage fates.

Transcription factor Yin Yang 1 (YY1) is a ubiquitously expressed factor that plays significant roles in normal biological processes including cell differentiation, proliferation, replication, DNA repair, and embryogenesis. Embryonic YY1 knockout results in peri-implantation lethality indicating its critical role during embryogenesis (Donohoe et al. 1999). YY1 derived its name based on its ability to both activate expression of some genes, while repressing expression of others (Hariharan et al. 1991; Park and Atchison 1991; Shi et al. 1991). YY1 can activate gene expression through recruitment of histone acetyltransferase (HAT) proteins and can positively regulate numerous genes (Bushmeyer et al. 1995; Galvin and Shi 1997). Moreover, YY1 can mediate long-distance enhancer-promoter DNA interactions perhaps through self-dimerization (Beagan et al. 2017; Weintraub et al. 2017), and is required for Ig locus contraction needed for immunoglobulin gene rearrangement and enhancer interactions for Ig class switch recombination (Ebert et al. 2011; Medvedovic et al. 2013; Mehra et al. 2016). Alternatively, YY1 can repress transcription of many genes by transient repression mechanisms involving recruitment of histone deacetyltransferase (HDAC) proteins (Galvin and Shi 1997). In addition, our laboratory found that YY1 can mediate Polycomb Group (PcG)-dependent repression that stably represses genes during development (Atchison et al. 2003; Wilkinson et al. 2006; Basu et al. 2010).

YY1 knockout early in the B lymphocyte lineage by action of Mb1-driven CRE results in developmental arrest at the pro-B cell stage (Liu et al. 2007). These arrested pro-B cells generate functional immunoglobulin heavy chain (IgH) gene V-D-J rearrangements that are skewed proximally within the IgH locus (Liu et al. 2007). Kleinman (Kleiman et al. 2016) found that deletion of YY1 at all stages of B cell development by action of CD19-CRE results in a negative impact at essentially all stages of B cell development, and we showed YY1 knockout driven by γ1-CRE activated in germinal center B cells, results in complete loss of these cells and absence of serum IgG1 (Banerjee et al. 2016). In addition to B cell lineage development, YY1 plays important roles in a multiplicity of developmental programs including T cells, intestinal stem cells, iNKT cells, cardiac and striated muscle, myeloid cells, mesoderm cells, erythroid cells, and extended pluripotent stem cells (Walowitz et al. 1998; Erkeland et al. 2003; Sucharov et al. 2003; Blattler et al. 2012; Perekatt et al. 2014; Ou et al. 2019; Perreault et al. 2020; Dong et al. 2022). The multiplicity of developmental programs impacted by YY1 suggests a common developmental mechanism among multiple lineages.

While analyzing transcripts in wild-type pro-B cells compared to YY1 knockout pro-B cells we observed reduced expression of several B lineage genes coincident with increased expression of some alternative hematopoietic lineage genes. This prompted us to explore whether YY1 knockout pro-B cells might be capable of differentiation into alternative lineages. We show here that YY1 knock-out pro-B cells acquire the capacity to develop into T lineage cells both *in vitro* and *in vivo*. These cells gain a T-like transcript profile while losing the B lineage transcript profile. Forced expression of YY1 ablates this lineage plasticity. Strikingly, single cell RNA-seq experiments indicate that during transition from the pro-B to T cell lineage, most cells transiently acquire transcript profiles representative of a multiplicity of hematopoietic lineages. Our results suggest that the dual activation and repression mechanisms of YY1 may mediate a universal lineage commitment mechanism that when disrupted enables cells to adopt numerous alternative fates.

## Results

### YY1-null pro-B cells show lineage plasticity in vitro and can develop into T lineage cells

Evaluation of published RNA seq data (Kleiman et al. 2016) revealed a number of B lineage-specific genes including Igll1, Vpreb1, Vpreb2, and CD79b, that showed reduced expression when comparing *yy1^f/f^* to *yy1^f/f^* Mb1-CRE (YY1-null) pro-B cells (Fig. S1A, left). However, the expression levels of key B-lineage regulatory factors such as Pax5, Ebf1, E2A, Foxo1, and Ikaros were unchanged (Figure S1A, right). Interestingly, expression of several hematopoietic lineage-specific genes was elevated (Figure S1B). Based on these unusual expression patterns, we questioned whether YY1-null pro-B cells, which should be committed to the B lineage pathway, exhibit lineage plasticity. To address this, we utilized a feeder cell line (OP9-DL4) that expresses the Notch ligand Delta-like 4 and promotes T cell development in the presence of IL7, SCF, and Flt3L (Mohtashami et al. 2013).

Pro-B cells from bone marrow (BM) were purified by fluorescence-activated cell sorting (FACS) from wild-type (*yy1^f/f^*) C57BL/6 mice, as well as from mice in which the *yy1* gene is deleted in pro-B cells by the action of Mb1CRE (*yy1^f/f^ Mb1-CRE*). Isolated pro-B cells were cultured on OP9-DL4 feeder cells in the presence of IL-7, Flt3L, and SCF (Mohtashami et al. 2010) (Figure 1A), and after three weeks, were harvested and analyzed for cell surface expression of T lineage marker proteins CD25 and Thy1.2. As anticipated, *yy1^f/f^* (wildtype) pro-B cells failed to express CD25 or Thy1.2 (Figure 1B left top and right panels) indicating B lineage commitment. However, over 90% of *yy1^f/f^ Mb1-CRE* (YY1-null) pro-B cells efficiently expressed T lineage marker proteins CD25 and Thy1.2 (Figure 1B left bottom and right panels). We further evaluated YY1-null pro-B cell derived CD25^+^ Thy1.2^+^ cells by FACS with anti-CD44 and anti-CD25 which distinguish double negative T lineage cells into 4 stages (DN1, DN2, DN3, and DN4). This evaluation showed that YY1-null pro-B cells predominantly developed *in vitro* into DN3 T lineage cells (Figure 1C). These cells contained IgH Vh7183-DJh rearrangements confirming their pro-B cell origin (Figure 1D, left panel), as well as T cell receptor (TCR) Vβ4 rearrangements consistent with their development into T-like cells (Figure 1D, right panel).

**Figure 1.**
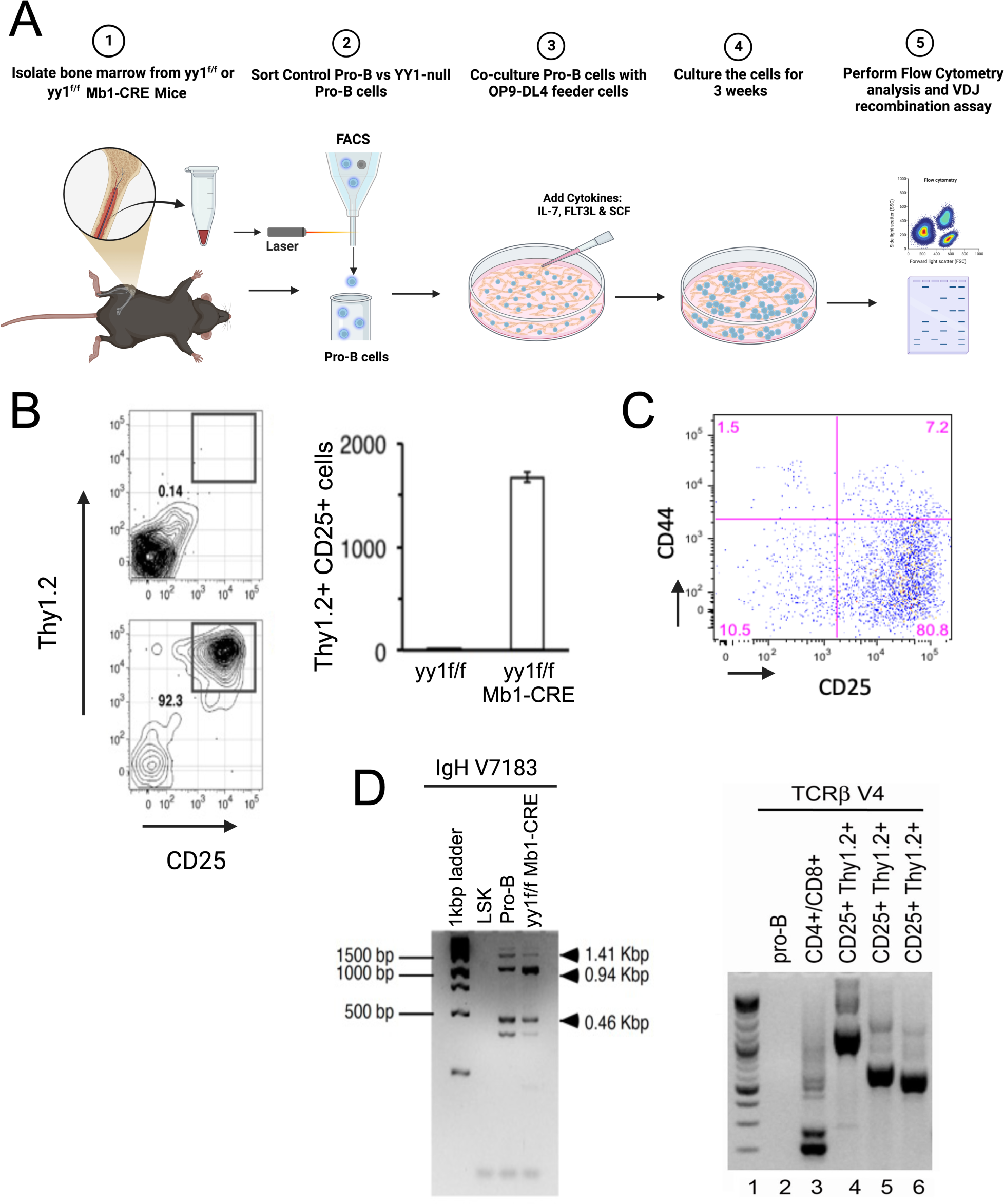
YY1 knockout pro-B cells can adopt T lineage properties *in vitro*. *(A)* Diagram of *in vitro* strategy for testing lineage plasticity of either wild-type (*yy1f/f*) or YY1 knockout (*yy1f/f Mb1-CRE*) pro-B cells. *(B)* Comparison of *yy1f/f* and *yy1f/f Mb 1CRE* pro-B cells grown on OP9-DL4 feeder cells in the presence of IL-7, Flt3L, and SCF for three weeks. Left panel: FACS plots of *yy1f/f* (top) compared to and *yy1f/f Mb 1CRE* (bottom) pro-B cells with antibodies that detect T lineage cells (Thy1.2 and CD25). Right panel: quantitation of cell numbers in each sample. *(C)* CD44 and CD25 antibody FACS profile of T lineage DN stages of *yy1f/f Mb1-CRE* pro-B cells grown on OP9-DL4 feeders for three weeks. *(D)* YY1 knockout pro-B cells grown on OP9-DL4 feeders for three weeks contain both IgH and TCRβ somatic rearrangements. Left panel: VDJ rearrangement assays for VH7183 rearrangements with LSK negative control and wild-type pro-B positive control. Right panel: TCRβ V4 rearrangements with pro-B negative control, CD4/CD8 T cell positive control and three samples of *yy1f/f Mb-1CRE* pro-B cells grown on OP9-DL4 feeder cells in the presence of IL-7, Flt3L, and SCF for three weeks.

### Ectopic expression of YY1 in YY1-null pro-B cells ablates their T lineage potential

To prove that loss of YY1 expression in pro-B cells was the cause of their acquired lineage plasticity, we performed YY1 complementation experiments (Figure 2A). Bone marrow cells isolated from *yy1^f/f^ Mb1-CRE* mice were transduced with either empty retroviral vector (MigR1) or with vector expressing the YY1 cDNA (MigR1-YY1) (Figure 2A). Transduced bone marrow cells were then injected into lethally irradiated C57BL/6 mice, and after 12 weeks mice were evaluated. We anticipated that YY1-null pro-B cells expressing the MigR1 vector alone would fail to support B lineage development past the pro-B cell stage as we and others previously demonstrated (Liu et al. 2007; Pan et al. 2013), but would be capable of T lineage development *in vitro* on OP9-DL4 feeders due to absence of YY1. Alternatively, YY1-null pro-B cells transduced with MigR1-YY1 would support B lineage development past the pro-B cell stage but would fail to develop into T lineage cells on DL4-OP9 feeders. As expected, in mice receiving MigR1 vector control, B lineage development was largely arrested at the pro-B cell stage due to absence of YY1 (Figure 2B top panel). B lineage cells failed to progress efficiently past the pro-B cell stage, yielding very few pre-B cells, immature B cells, or recirculating B cells (Figure 2B, top panel). However, YY1-null pro-B cells isolated from the BM of these mice were capable of generating CD25^+^ Thy1.2^+^ T lineage cells on OP9-DL4 feeders consistent with their lack of YY1 (Figure 2B, bottom panel). On the contrary, mice injected with cells transduced with MigR1-YY1 fully recovered B lineage development due to the presence of retrovirally expressed YY1, efficiently generating pre-B, immature B, and recirculating B cells (Figure 2C, top panel). Moreover, pro-B cells isolated from these mice were incapable of generating cells with T lineage surface markers CD25^+^ Thy1.2+ on OP9-DL4 feeders (Figure 2C bottom panel). Thus, we conclude that absence of YY1 is the regulating factor enabling lineage plasticity of normally committed pro-B cells.

**Figure 2.**
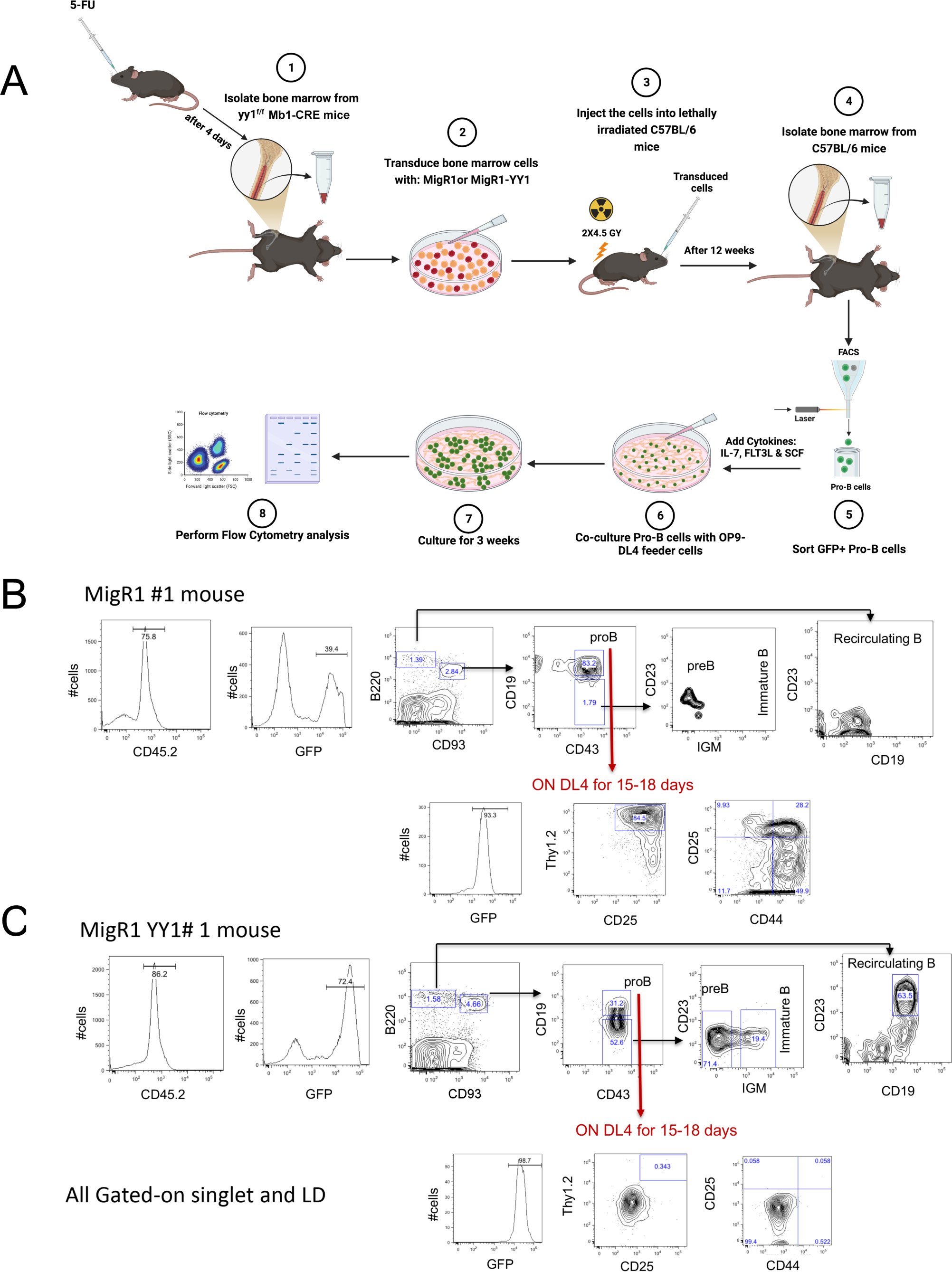
YY1-complementation ablates lineage plasticity and restores B lineage commitment. *(A)* Diagram of the experimental strategy. *(B)* Empty retroviral vector MigR1 does not rescue B lineage arrest caused by YY1 knockout and allows T lineage plasticity of *yy1* knockout pro-B cells. Bone marrow cells from *yy1f/f Mb-1CRE* mice were isolated, transduced with MigR1 empty vector, and injected into lethally irradiated mice. Twelve weeks later, GFP+ pro-B cells were isolated and placed on OP9-DL4 feeder cells for 3 weeks. FACS analyses showed development of pro-B cells but not of pre-B or recirculating B lineage cells (top panel). These cells generated T lineage cells on OP9-DL4 feeders as assessed by FACS with anti-Thy1.2, CD25, and CD44 antibodies yielding patterns similar to DN2 and DN3 cells (bottom panel). *(C)* Retroviral provision of YY1 restores B lineage development but ablates lineage plasticity to the T cell lineage. Bone marrow cells from *yy1f/f Mb-1CRE* mice were isolated, transduced with YY1-expressing vector MigR1-YY1, and injected into lethally irradiated mice. Twelve weeks later, GFP^+^ pro-B cells were isolated and placed on OP9-DL4 feeder cells for 3 weeks. YY1 expression completely rescued B lineage development by formation of pro-B, pre-B, immature B, and recirculating B lineage cells *in vivo* (top panel), but these cells failed to generate T lineage cells on OP9-DL4 feeder cells (bottom panel).

### YY1-null pro-B cell-derived T-like cells exhibit transcript profiles resembling wild-type thymic T cells

To evaluate the similarity of the T-like cells developed from YY1-null pro-B cells to wild-type thymic T cells isolated from mice, we performed RNA-seq studies with RNA prepared from 5 independent experiments. RNA-seq profiles of YY1-null pro-B cells developed into T-like cells (T) and control wild-type thymic T cells (C), were evaluated and compared with already available RNA-seq profiles of wild type pro-B cells (B), and YY1-null pro-B cells (Y) (Kleiman et al. 2016). Principal component analyses (PCA) of all samples indicated that the YY1-null pro-B–derived T-like lineage cells were very distinct from wild-type and YY1-null pro-B cells, and clustered adjacent to wild-type thymocytes (Figure 3A, left panel). Transcript correlation distances by cluster dendrogram also showed YY1-null derived T-like cells were much more similar to T cells than to pro-B cells (Figure 3A, right panel).

**Figure 3.**
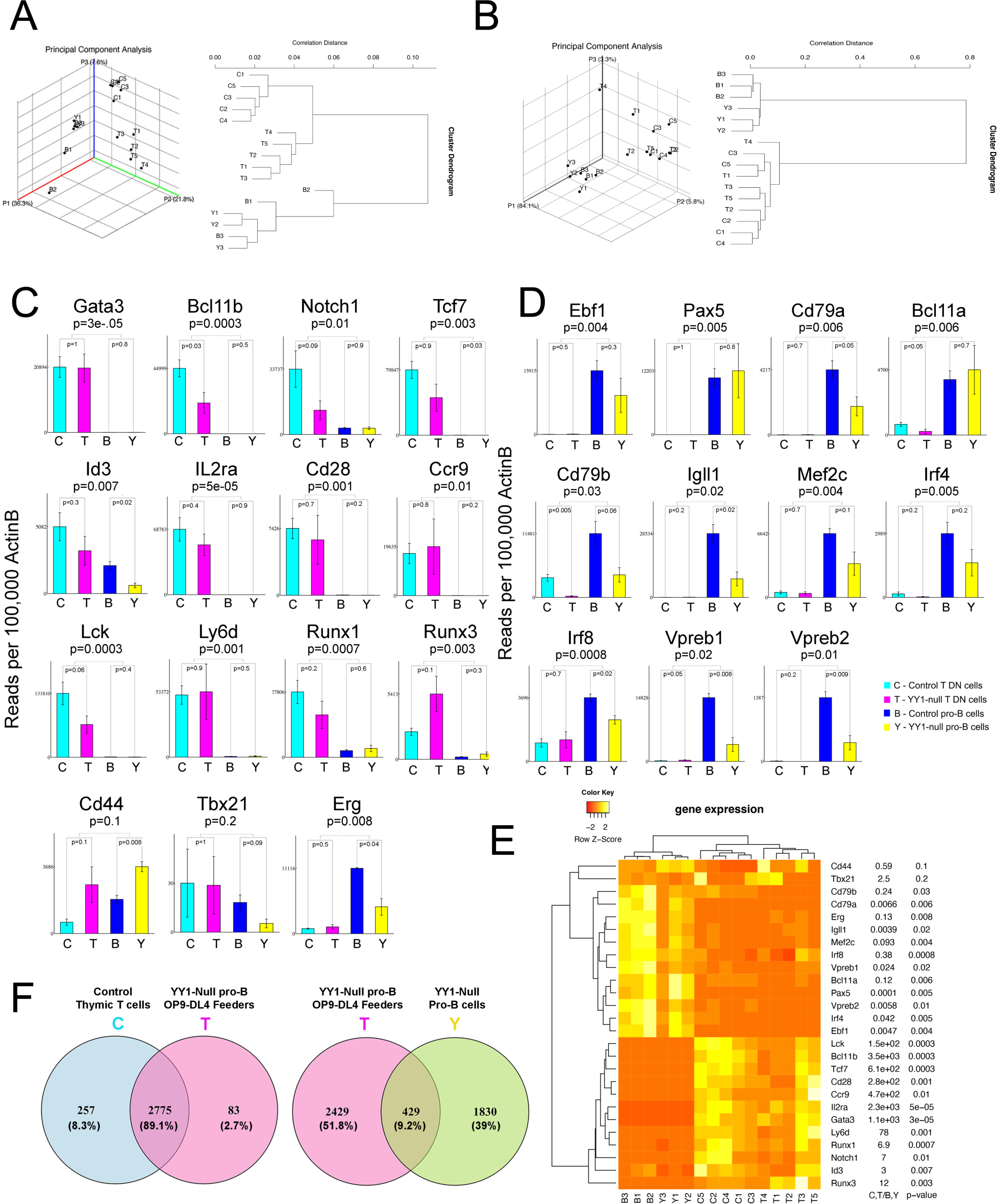
YY1 knockout pro-B cells grown on OP9-DL4 feeders for three weeks gain a T lineage transcript profile and ablate their B lineage profile. *(A)* 3-dimensional Principal Component Analyses (PCA) and dendograms of the RNA-seq data from Control thymic T cells (C1-C5), T cells developed from YY1-null pro-B cells (T1-T5), normal pro-B cells (B1-B3) and YY1-null pro-B cells (Y1-Y3). *(B)* 3-dimensional PCA and dendograms of RNA-seq data from 15 select T cell and 11 select B cell genes. *(C)* Gain of T lineage RNA-seq profiles of *yy1f/f Mb1CRE* pro-B cells grown of OP9-DL4 feeders for three weeks. RNA-seq expression levels are shown for select T lineage genes using RNA isolated from YY1-knockout pro-B cells (Y), control pro-B cells (B), YY1-knockout pro-B cells grown on OP9-DL4 feeders for three weeks (T), and control thymic T cells (C). *(D)* Loss of B lineage gene expression in *yy1f/f Mb1CRE* pro-B cells grown of OP9-DL4 feeders for three weeks. Gene expression profiles are shown for select B lineage genes using the same RNA-seq samples as in panel C. *(E)* Heat map expression profiles of the data in panels C and D. *(F)* Overlap in gene expression profiles in each sample. (Left) Control thymic T cell transcripts show extensive overlap in RNA-seq profile (89.1%) with *yy1f/fMb1CRE* pro-B cells grown on OP9-DL4 feeders (T). (Right) On the contrary *yy1f/fMb1CRE* pro-B cells grown on OP9-DL4 feeders (T) show expression of only 9.2% genes expressed in their original YY1-null pro-B cell identify (Y).

We evaluated a set of 15 genes representative of the T cell lineage, and 11 genes of the B cell lineage to determine if T-like cells developed from YY1-null pro-B cells gained a T lineage gene expression profile, while simultaneously extinguishing the B lineage gene expression profile. PCA and cluster dendrograms of expression patterns of the 15 T lineage and 11 B lineage genes indicated that YY1-null pro-B cell derived T-like cells clustered with wild-type thymic T lineage cells and were distinct from wild-type pro-B or YY1-null pro-B cells (Figure 3B). Transcript profiles of each of the 15 key T lineage genes Gata3, Bcl11b, Notch1, Tcf7, Id3, Il2ra, Cd28, Ccr9, Lck, Ly6d, Runx1, Runx3, and Erg all showed significantly increased expression in T-like cells developed from YY1-null pro-B cells (T), and closely matched the expression pattern in control wild-type thymic T cells (Figure 3C, C and T data samples). These genes were nearly silent in wild-type pro-B cells (B) as well as YY1 knockout pro-B cells (Y) (Figure 3C, B and Y samples). Only expression of Cd44 and Tbx1 showed insignificant differences between wild-type T cells and YY1-null pro-B cell-derived T-like cells compared to pro-B cell samples (Figure 3C). Similarly, we found that critical B lineage genes Ebf1, Pax5, Cd79a, Bcl11a, Cd79b, Igll1, Mef2c, Irf4, Irf8, Vpreb1, and Vpreb 2 all significantly dropped in expression in T-like cells derived from YY1-null pro-B cells, with expression patterns resembling wild type thymic T cells (Figure 3D). Heatmap representation of the 15 T lineage and 11 B lineage genes in each RNA-seq sample is shown in Figure 3E.

We also evaluated genes that were differentially expressed between our RNA-seq samples. A comparative analysis of data sets of YY1-null pro-B developed into T-like cells (T) versus wild-type thymic T cells (C) or YY1-null pro-B cells (Y) identified a total of 3116 and 2552 significant differentially expressed genes (DEGs) respectively, with log2-fold change in the range of ≥ +1.0 to ≤ −1.0. Out of 3116 DEGs, 692 were upregulated (log2FC ≥ +1.0 with p-value ≤ 0.05) and 2424 were downregulated (log2FC ≤ −1.0 with p-value ≤ 0.05) in YY1-null pro-B cells developed into T-like cells (T) vs wild-type thymic T cells (C). In wild type pro-B (B) vs YY1-null pro-B cells (Y) data, 993 genes were upregulated (log2FC ≥ +1.0 with p-value ≤ 0.05) and 1559 were downregulated (log2FC ≤ −1.0 with p-value ≤ 0.05) out of 2552 DEGs.

Expression of 89.1% of differentially expressed genes (2775 genes) were shared between wild-type thymic T cells and YY1 knock-out pro-B cells developed into T-like cells (Figure 3F, left panel). A very small fraction of genes (83 genes; 2.7%) were uniquely expressed in YY1-null pro-B cell-derived T-like cells, and 257 genes (8.3%) were unique to control thymic T cells (Figure 3F, left panel). Gene ontology (GO) analyses showed the unique genes in control T lineage cells compared to YY1 knockout derived T-like cells grouped into a variety of pathways that were not specific to T lineage cells (Figure S1C, left panel, red lettering). Similarly, the pathways uniquely expressed in YY1-null pro-B cells developed into T-like cells showed mainly non-T lineage-specific pathways (Figure S1C, right panel, red lettering). On the contrary, T-like cells developed from YY1-null pro-B cells showed extensive GO differences with YY1-null pro-B cells (Figure S1D left panel). Only 9.2% of transcripts overlapped between these samples, whereas 90.8% of transcripts were unique (Figure 3F, right panel). GO analyses showed that T-like cells developed from YY1-null pro-B cells expressed pathways enriched in T lineage cells (Figure S1D, left panel, blue lettering). On the contrary, as expected, YY1-null pro-B cells expressed pathways indicative of their B cell lineage (Figure S1D, right panel, green lettering). Intriguingly some T lineage and other lineage pathways were also expressed in in YY1-null pro-B cells (Figure S1C, right panel blue lettering). These differences are consistent with YY1 knockout pro-B cells perhaps being capable of generating other lineages in addition to T cells.

### YY1-null pro-B cells can differentiate into T-like cells in vivo

Our studies using OP9-DL4 feeder cells clearly showed that YY1-null pro-B cells can develop into cells that closely resemble thymic T lineage cells. We questioned whether YY1-null pro-B cells could differentiate into T-like cells *in vivo*, and perhaps past the DN T cell stages. To test this, we isolated by FACS wild-type pro-B cells from *yy1^f/f^* mice and YY1-null pro-B cells from *yy1^f/f^ mb1-CRE* mice and injected these cells into sub-lethally irradiated Rag1^-/-^ mice which fail to develop either mature T lineage or B lineage cells (Figure 4A). Any developing T-like cells therefore, would be due to the injected pro-B cells. After 7 months Rag1^-/-^ mice injected with wild-type pro-B cells failed to develop detectable thymuses (Figure 4B, top panel). On the contrary, 3 out of 15 Rag1^-/-^ mice injected with YY1-null pro-B cells developed large thymuses, with the remaining developing smaller rudimentary thymuses (Figure 4B, bottom panel). Thymocytes or splenocytes isolated from these mice with the larger thymuses showed the acquisition of T-like double positive (DP), and single positive CD4 and CD8 cells, respectively (Figure 4C), and their new T-like phenotype was confirmed in splenic T cells by the expression of TCRβ protein on the cell surface (Figure 4D). Consistent with their pro-B cell origin, these cells also possessed IgH VhQ52 rearrangements (Figure 4E)

**Figure 4.**
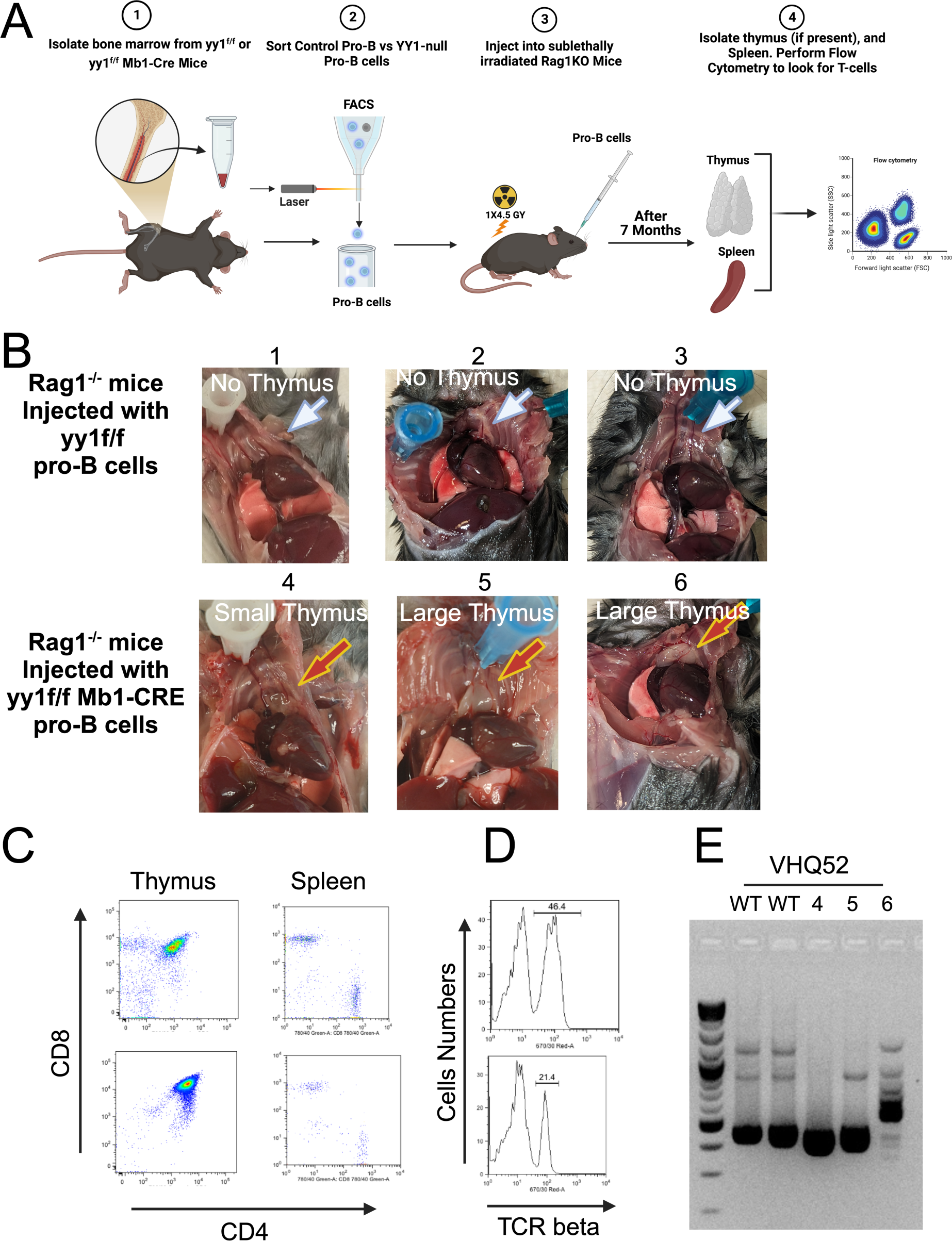
YY1 knockout pro-B cells can develop into T-like cells *in vivo*. *(A)* Diagram showing experimental strategy. *(B)* Sub-lethally irradiated Rag1^-/-^ mice were injected with either control pro-B cells (*yy1f/f*), or YY1 knockout pro-B cells (*yy1f/f Mb1CRE*). Seven months later mice were sacrificed and assessed for T lineage development. Pictures are shown of mouse thymuses from either control Rag1^-/-^ mice injected with wild-type (*yy1f/f*) pro-B cells (top panels) or injected with YY1 knockout pro-B cells (*yy1f/f Mb1CRE*; bottom panels). *(C)* FACS plots of thymic DP, CD4 and CD8 T lineage development (left panel) of two mice injected with *yy1f/f Mb1CRE* pro-B cells, and CD4+ CD8+ expression of splenic T cells of 2 mice injected with *yy1f/f Mb1-CRE* pro-B cells (right panel). *(D)* Splenic T lineage cells developed *in vivo* from *yy1f/f Mb1CRE* pro-B cells express TCRβ. *(E)* Two of three samples of T lineage cells developed *in vivo* from *yy1f/f Mb1CRE* pro-B cells exhibit IgH gene rearrangements (samples 5 and 6).

### Alternative hematopoietic lineage genes are activated during transition from the pro-B to T cell lineage

The transformation of YY1-null pro-B cells into T-like cells on OP9-DL4 feeders required approximately 3 weeks to generate a population that consisted of over 90% CD25^+^ Thy1^+^ cells. We reasoned that harvesting cells mid-way during this process and subjecting them to single-cell RNA-seq (sc-RNA-seq) would enable us to decipher key mechanistic details of this alternative lineage differentiation process. Therefore, we isolated pro-B cells from *yy1f/f* and *yy1f/f mb1CRE* mice and cultured them on OP9-DL4 cells with IL7, Flt3L, and SCF until 3 to 5% of the *yy1f/f mb1CRE* pro-B population had become CD25^+^ Thy1^+^ (as expected, cells from wild-type *yy1f/f* mice remained negative). Cells cultured on OP9-DL4 feeders were harvested and subjected to scRNA-seq in parallel with pro-B cells isolated directly from *yy1f/f* and *yy1f/f mb1CRE* mice.

Merged scRNA-seq data from duplicate samples of the four conditions (pro-B cells from *yy1f/f* and *yy1f/f Mb1CRE* mice, plus the two pro-B samples on OP9-DL4 feeders) generated 22 Seurat UMAP clusters (Figure S2A). Separating the overlapping merged UMAPs into the individual sample UMAPs showed very similar patterns comparing *yy1f/f* and *yy1f/f Mb1CRE* pro-B cells directly from mice, as well as *yy1f/f* pro-B cells cultured on OP9-DL4 feeders (Figure 5A, panels 2-4). However, the UMAP of the *yy1f/f Mb1CRE* pro-B cells cultured on OP9-DL4 feeders showed considerable differences with reductions of cell numbers in clusters 0, 1, 2, 4, 14, and 21, but substantial gains in clusters 3, 7, 8, 13, 18, and 20 (Figure 5A, panel 1). We used the SingleR package in R Studio for cell annotation using the Immgen database to define the different cell types present in each cluster. This analysis annotated nearly all the cells in the *yy1f/f* and *yy1f/f Mb1CRE* pro-B cells directly from mice, as well as with the *yy1f/f* samples grown on OP9-DL4 feeders as B lineage cells (Figure 5B, panels 2-4; pink-orange color). On the contrary, nearly all the cells in the *yy1f/f Mb1CRE* sample grown on OP9-DL4 feeders had lost their B lineage phenotype and expressed transcripts specific for numerous distinct hematopoietic cell types indicated by multiple non-B cell lineage colors. (Figure 5B, panel 1). This panel is expanded in Figure 5C with cell types identified by the SingleR program overlaid on the figure. Cell types annotated in this sample included T, Tψ8, ILC, NKT, NK, dendritic, monocyte, basophil, macrophage, mast, neutrophil, and stem cell phenotypes (Figure 5C). The 3-5% of cells identified as T lineage when the population was harvested for scRNA-seq analysis are shown by the arrow in Figure 5C.

**Figure 5.**
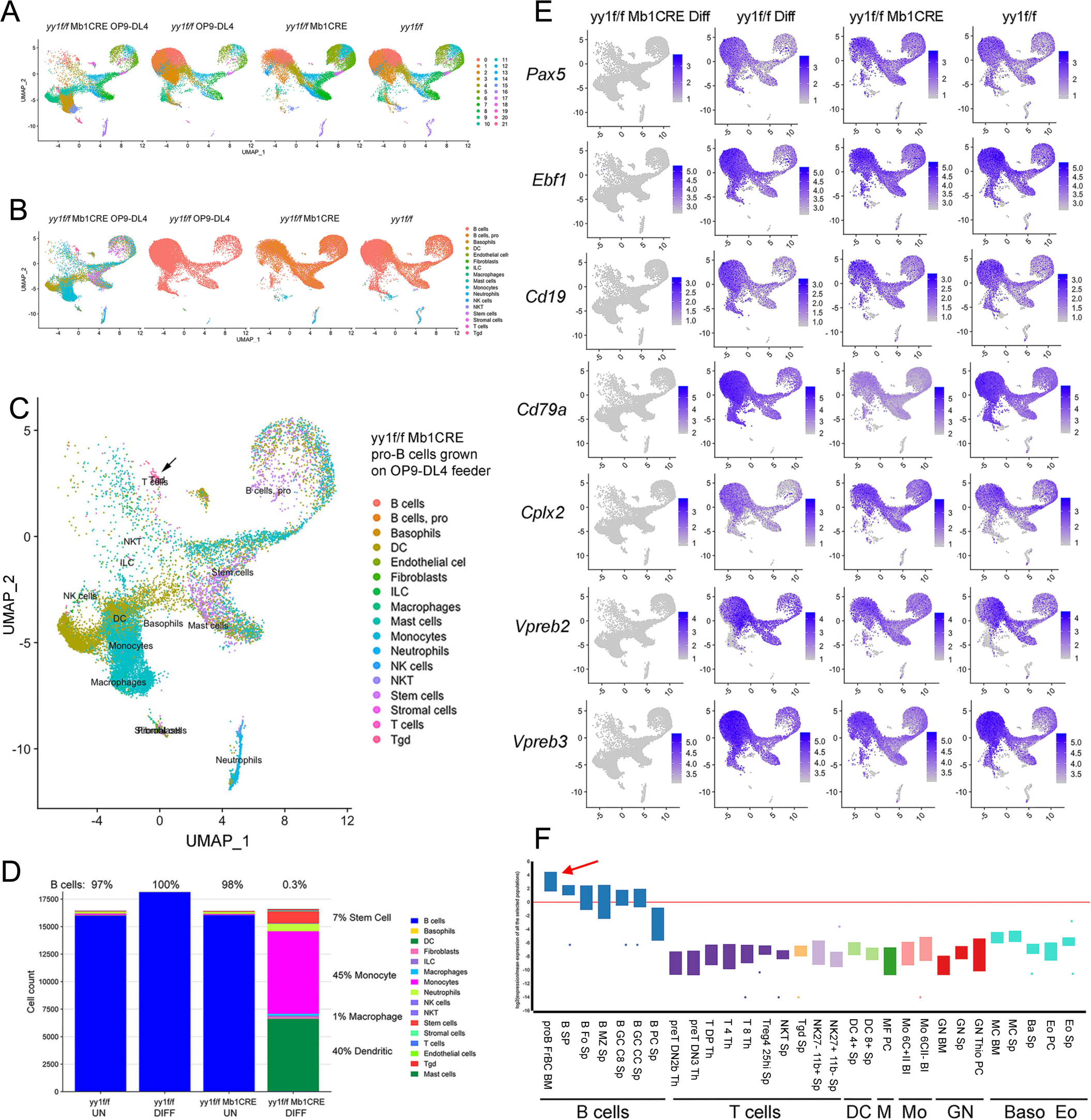
*yy1f/f Mb1CRE* pro-B cells grown on OP9-DL4 feeders for 2 weeks develop into cells expressing a multiplicity of hematopoietic lineage transcripts, and extinguish their B lineage transcript profile. Pro-B cells from *yy1f/f* or *yy1f/f Mb1CRE* mice were either subjected to scRNA-seq immediately or grown on OP9-DL4 feeders for 2 weeks when 3-5% of cells from the yy1f/f Mb1-CRE sample developed a Thy1.2^+^ CD25^+^ phenotype. *(A)* Patterns of the 22 Seurat clusters in each sample. *(B)* Assignment of cell types in each sample using the SingleR program. Cell identity key is in the right panel. *(C)* Pattern of the clusters and cell types in the *yy1f/f Mb1CRE* sample grown on OP9-DL4 feeders. Cell types identified by SingleR are indicated within the figure. The arrow points to the location of cells defined as either T or Tψ/8 lineage. *(D)* Cell type percentages in each sample. The percentage of each sample that represents B lineage cells is shown above each column. The percentage of cells in the *yy1/f/f Mb1CRE* sample grown on OP9-DL4 feeders that are identified as either stem cell, monocyte, macrophage, or dendritic cells is indicated, and cell identities are indicated in the color key on the right. *(E)* UMAP profiles are shown for B lineage genes Pax5, Ebf1, Cd19, Cd79a, Cplx2, Vpreb2, and Vpreb3 from pro-B cells isolated from yy1f/f and yy1f/f Mb1-CRE mice, or the same cells grown on OP9-DL4 feeders for 2 weeks until 3-5% of the cells from yy1f/f Mb1-CRE mice were Thy1.2^+^ CD25^+^. Gene names are on the left and above each column are the cell type. *yy1f/f Mb1CRE* Diff are YY1 knockout pro-B cells developed until 3-5% of the pro-B cells developed a Thy1.2^+^ CD25^+^ phenotype, *yy1f/f* Diff are wild-type pro-B cells incubated on feeders for the same period of time (2 weeks), and *yy1f/f Mb1CRE* and *yy1f*/f are pro-B cells directly isolated from mice and subjected to scRNA-seq. *(F)* Box plot charts of the genes listed in panel E for expression level in various B, T, Dendritic (DC), Macrophage (M), Monocyte (Mo), Granulocyte (GN), basophil (Baso), or eosinophil (Eo) cell types. The red arrow points to pro-B fraction B/C cells.

Percentage of various cell types in the entire population are shown in Figure 5D. The percentage of cells assigned by SingleR that were identified as B cells in the *yy1f/f* and *yy1f/f Mb1CRE* cells from mice, and the *yy1f/f* cells on OP9-DL4 feeders ranged from 97% to 100% (Figure 5D). On the contrary, only 0.3% of the cells (48 cells out of over 16,000) were annotated as B lineage cells in the *yy1f/f Mb1CRE* pro-B cells grown on OP9-DL4 feeders (Figure 5D). Forty-five percent of the cells were identified as monocytes, 40% as dendritic cells, 7% as stem cells, 4% as neutrophils, 1% as macrophages, and 1% as ILC cells (Figure 5D).

The remarkable difference of the *yy1f/f Mb1CRE* pro-B cells on OP9-DL4 feeders compared to the other three samples indicate that the B lineage transcript profile in these cells is extinguished, and genes expressed in alternative lineages are elevated. Therefore, we set out to evaluate expression of genes closely associated with specific hematopoietic lineages.

### Extinguished B lineage expression patterns

We evaluated expression of key genes indicative of the B cell lineage. As expected, we found pro-B cells directly isolated from *yy1f/f* and *yy1f/f mb1CRE* mice, as well as control *yy1f/f p*ro-B cells grown on OP9-DL4 feeders highly expressed critical B lineage genes Pax5, Ebf1, Cd19, Cd79a, Cplx2, Vpreb2, and Vpreb3 throughout their UMAP profiles (Figure 5E, UMAP panels 2-4). Quite strikingly however, expression of each of these B lineage genes was completely extinguished in the *yy1f/f mb1CRE* pro-B cells grown on OP9-DL4 feeders (Fig. 5E, UMAP panel 1). Box plot graphs from the Immgen MyGeneSet program against various B lineage cells (pro-B fraction B and C, Splenic B, follicular B, marginal zone B, germinal center centroblasts, germinal center centrocytes, and splenic plasma cells) indicated high correlation of these transcripts (Pax5, Ebf1, Cd19, Cd79a, Cplx2, VpreB2, and VpreB3) with the B lineage, with highest expression in pro-B cells (Figure 5F, red arrow). In contrast, there was very low correlation of these genes in numerous T lineage, dendritic, monocyte, granulocyte, basophil, and eosinophil cell types (Figure 5F). The dramatic loss of B lineage gene transcripts in the *yy1f/f Mb1-CRE* pro-B cells grown on OP9-DL4 feeders (Figure 5E, panel 1) is consistent with the SingleR cell-type annotation program indicating only 0.3% of cells retained a B lineage transcript profile (Figure 5C and D).

### Gain of T cell gene expression

The OP9-DL4 *in vitro* differentiation system is designed to generate T cells from precursor cells (Mohtashami et al. 2013). In our scRNA-seq experiments, we halted the culture system when only a small fraction of cells (less than 5%) had generated a Thy1.2^+^ CD25^+^ surface phenotype. If incubated to completion (3 weeks) we showed that most cells developed to the DN3 T-like cells (Figure 1C). Therefore, we set out to determine the identities of genes that are enriched in DN3 cells compared to pro-B cells, then to evaluate expression of those genes in our scRNA-seq data. First, we used the Immgen MyGeneset program to compare traditional RNA-seq data from pro-B fraction B/C cells with DN3 cells. The top 25 genes enriched in DN3 cells compared to pro-B cells are shown in Table S1. This comparison identified *Cd3g, Trbc1, Bcl11b, Cd3d, Tcf7*, and *Trbc2* within the top 10 enriched genes, while *Trdc* and *Trgv2* were in the top 25 enriched genes. *Thy1* was the 47^th^ most enriched gene. We evaluated expression of each of these genes in UMAPs of our four scRNA-seq samples (*yy1f/f Mb1CRE* pro-B cells differentiated on OP9-DL4 feeders until 5% of cells were *Thy1.2^+^ CD25*^+^; *yy1f/f* pro-B cells differentiated for the same time on OP9-DL4 feeders; *yy1f/f Mb1CRE* pro-B cells and *yy1f/f* pro-B cells both isolated directly from mice). High gene expression levels were observed for each of the key DN3 T lineage genes in the *yy1f/f Mb1CRE* differentiated sample within the small cluster of cells identified as either T cells or Tψ8 cells by SingleR (Figure 6A red circle and Figure 6B red arrows, respectively). Scattered expression was observed in other cells throughout the UMAPs in the differentiated *yy1f/f Mb1CRE* sample in Figure 6B consistent with nearly all cells ultimately adopting a T lineage phenotype if incubated longer (3 weeks). Either no expression, or very low expression of DN3 T lineage genes was observed in each of the control scRNA-seq samples (undifferentiated *yy1f/f Mb1CRE* pro-B cells, or the *yy1f/f* pro-B cells either directly from mice or incubated on OP9-DL4 feeders; Figure 6B, panels 2-4). We also evaluated expression of *Cd3g, Trbc1, Bcl11b, Cd3d, Tcf7, Trdc, Trgv2, and Thy1* in violin plots of each scRNA-seq cell cluster from the differentiated *yy1f/f Mb1CRE* sample. Consistent with our data in Fig. 6B, highest level expression was observed in cells from Seurat cluster 0 which SingleR identified as T cells (Figure 6C). Immgen MyGeneSet box plots confirmed expression of our DN3 enriched genes are highly specific for various T cell subsets (Figure 6D, red arrow).

**Figure 6.**
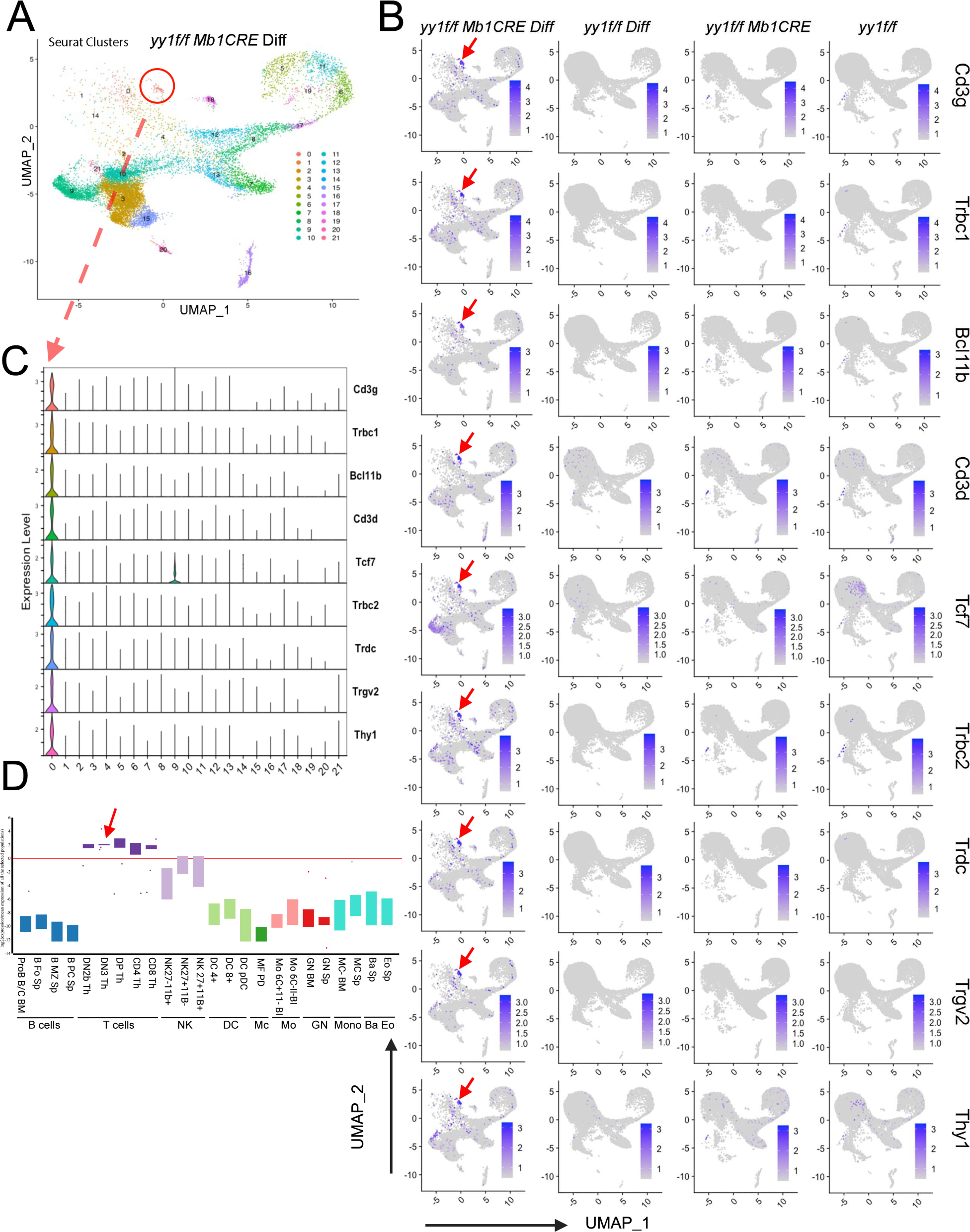
T lineage genes are expressed in the small cell cluster identified as T cells. *(A)* UMAP profile of *yy1f/f Mb1CRE* pro-B cells grown on OP9-DL4 feeders for two weeks until 3-5% of cells were Thy1.2^+^ CD25+. The red circle identifies the small cluster of cells identified by singleR as T lineage cells in the *yy1f/f Mb1CRE* sample. *(B)* UMAP gene expression profiles of key T cell lineage genes listed on the right. The red arrows show the location of the small cell cluster identified by SingleR as T lineage cells. *(C)* Violin plots of RNA expression level of each T lineage gene in each Seurat cluster. The dashed red arrow shows that high expression of T lineage genes matches the small cluster of cells in panel A identified as T cells (red circle). *(D)* Box plot charts of the genes listed in panel B for expression level in various B, T, NK, Dendritic (DC), Macrophage (Mc), Granulocyte (GN), Monocyte (Mo), basophil (Baso), or eosinophil (Eo) cell types.

### Gain of Dendritic cell-like expression patterns

Forty percent of *yy1f/f Mb1CRE* pro-B cells grown on OP9 DL4 feeders were identified as dendritic cells in our scRNA-seq experiments (Figure 5C and D). Using the Immgen MyGeneset program with the Immgen RNA-seq databases of DC4+ and DC8+ splenic dendritic cells compared to pro-B fraction B/C data, we identified Clec9a, Tlr11, Gm6377, Itgax, Slamf8, Gpr141b, Gpr35, Gpr34, Xcr1, Havcr2, Ifi205, Il1b, Cd83, Apol7c, and Cacnb3 genes as 15 of the top 25 genes highly enriched in DC4+ and DC8+ dendritic cells (Table S1). SingleR identified Seurat clusters 9 and 10 in our *yy1f/f Mb1CRE* differentiated sample as dendritic cells (see red ovals in Figure 7A). Indeed, dendritic genes Apol7c, Cacnb3, Clec9a, Tlr11, Gpr141b, and Xcr1 were highly expressed in these clusters in *yy1f/f Mb1CRE* pro-B cells differentiated on OP9-DL4 feeders for 14 days (Figure 7B, panel 1, red arrows). Essentially no expression was observed in the control panels (Figure 7B, panels 2-4). The other DC4+ and DC8+ dendritic genes (Gpr35, Gpr34, Gm6377, Itgax, Slamf8, Havcr2, Ifi205, Il1b, and Cd83) were also highly expressed in Seurat clusters 9 or 10, (Figure S2B, red oval). These genes were also expressed in the region identified as monocytic cells (clusters 3 and 15) consistent with the ability of monocytes to develop into dendritic cells (Figure S2B and C, blue circle and arrows, respectively). While the dendritic genes tested were highly expressed in YY1 knockout pro-B cells on OP9-DL4 feeders, they were silent or poorly expressed in the three control cell types in Figure 7B and Figure S2C, panels 2-4 indicating that YY1 is critical for repressing these dendritic genes in the OP9 DL4 differentiation system.

**Figure 7.**
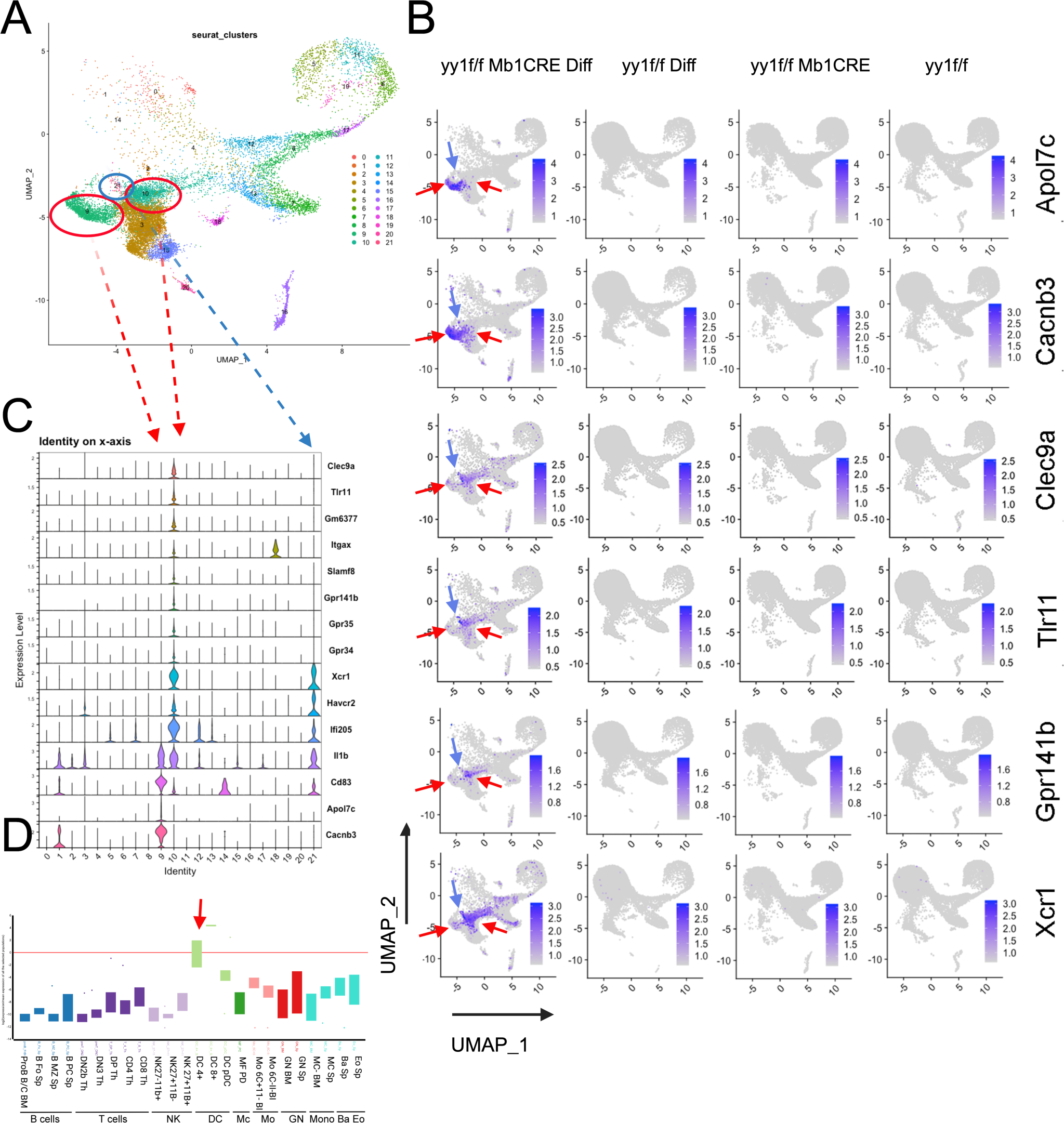
Elevated DC4+ and DC8+ dendritic lineage gene expression in clusters identified by SingleR as dendritic. *(A)* UMAP profile of yy1f/f Mb1-CRE pro-B cells grown on OP9-DL4 feeders for 2 weeks until 3-5% of cells were Thy1.2+ CD25+. Cell clusters 9 and 10 identified by SingleR as dendritic cells are indicated by red ovals. Cluster 21 also expressing dendritic genes is indicated by the blue oval. *(B)* UMAP profiles of gene expression for dendritic genes Apol7c, Cacnb3, Clec9a, Tlr11, Gpr141b, and Xcr1 show high level expression in clusters 9 and/or 10 (red arrows). Expression of some dendritic genes in cluster 21 is shown by the blue arrows. *(C)* Violin plots of RNA expression show that expression of the 6 dendritic genes in panel B, as well as 9 other dendritic genes match the dendritic clusters 9, 10 (red dashed arrows) and cluster 21 (blue dashed arrows). *(D)* Box plot charts of the genes in Panel B for expression level in various B, T, NK, Dendritic (DC), Macrophage (Mc), Granulocyte (GN), Monocyte (Mo), Basophil (Baso), or Eosinophil (Eo) cell types show strong specificity for DC4+ and DC8+ dendritic cells (red arrow).

Violin plots confirmed expression of the 15 DC4+ or DC8+ dendritic genes in clusters 9 and 10 (Figure 7C). Some genes (Xcr1, Vavcr2, Ifi205, Il1b, and Cd83) were also expressed highly in cluster 21 directly adjacent to dendritic clusters 9 and 10 (Figure 7A blue circle and 7C dashed blue arrow). Immgen MyGeneSet box plots confirmed that expression of these genes is specific for DC4+ and DC8+ dendritic cells (Figure 7D, red arrow). Thus, YY1 knock-out pro-B cells on OP9-DL4 feeders yield a transient phenotype consistent with dendritic cells.

We also evaluated transcripts representative of the DCpDC dendritic cell type. Immgen MyGeneSet analyses of RNA-seq data from DCpDC dendritic cells compared to pro-B fraction B/C cells, identified Ccr9, Siglech, Pacsin1, Klk1, Gm5547, Press30, Lag3, Pltp, and Runx2 as within the top 25 DCpDC expressed genes compared to pro-B cells (Table S1). Expression of these genes was highly elevated in cluster 18 in *yy1f/f Mb1CRE* samples grown on OP9-DL4 feeders, with very low-level expression in each of the control scRNA-seq samples (Figure S3A, red circle and S3B, panel 1, red arrows). In some cases (Pltp, Prss30, Gm5547), high level expression was also observed in cluster 3 (monocytic cells) (Figure S3B, panel 1, blue arrows), consistent with monocytes giving rise to dendritic cells. Violin plots confirmed elevated expression of DCpDC genes in cluster 18 (Figure S3C red dashed line), and the specificity of these genes for DCpDC cells is also confirmed by box plots (Figure S5D, red arrow).

### Gain of Monocyte cell-like expression patterns

SingleR identified the largest fraction of cells in the *yy1f/f Mb1CRE* pro-B cells grown on OP9-DL4 feeders as monocytes residing in clusters 3 and 15 (45% of cells, Figure 5C and D). To confirm this phenotype, we used the Immgen public database of transcripts expressed in monocyte populations (Monocytes 6C^-^ II^-^ BI and 6C^+^ II^-^ BI) compared to those in pro-B cells (fraction B/C). We identified genes Klra2, Arhgef37, Slfn1, Cd300ld, Ptpro, Gda, Gm9733, Ccl9, and Clec4a3 as being within the top 25 genes expressed in monocytes relative to pro-B cells (Table S1). These genes showed high transcript expression in clusters 3 and 15 (Figure 8A red circles and Figure 8B, panel 1 red arrows). These genes were poorly expressed in the control scRNA-seq UMAPS (Figure 8B, panels 2-4). Violin plots also showed very high expression of these genes in clusters 3 and 15 (Figure 8C, red dashed arrows). Immgen MyGeneSet box plots confirmed the strong correlation of Klra2, Arhgef37, Slfn1, Cd300ld, Ptpro, Gda, Gm9733, Ccl9, and Clec4a3 genes with the monocyte lineage (Figure 8D, red arrow). These data show that the cells identified as monocytes in Figure 6C, indeed express genes highly enriched in monocytic cells.

**Figure 8.**
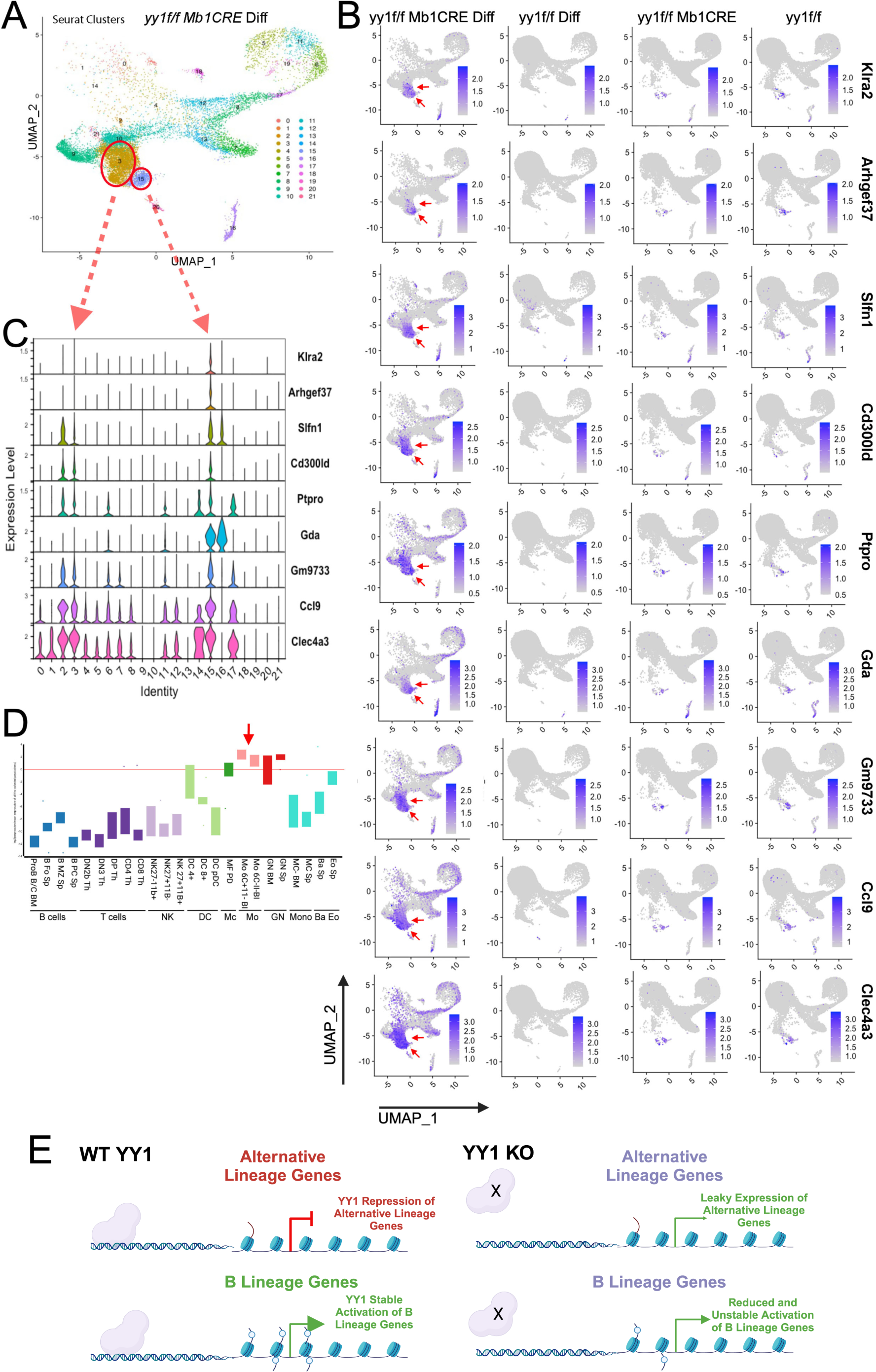
Elevated expression of monocyte genes in *yy1f/f Mb1CRE* cells on OP9 DL4 feeders, and model of YY1 control of B lineage commitment. *(A)* UMAP profile of yy1f/f Mb1-Cre pro-B cells from OP9-DL4 feeders grown for 2 weeks until 3-5% of the cells were Thy1.2+ Cd25+. The position of clusters identified as monocytic cells (clusters 3 and 15) are indicated by the red oval and circle. *(B)* UMAP profiles of gene expression of Klra2, Arhgef37, Slfn1, Cd300ld, Ptpro, Gda, Gm9733, Ccl9, and Clec4a3 show high level expression in clusters 3 and 15 (red arrows). *(C)* Violin plots also showed high expression of these genes in clusters 3 and 15 (red dashed arrows). *(D)* Immgen box blots show the genes presented in panels B and C are enriched in for expression in monocytes (red arrow). *(E)* Model of YY1 control of B lineage commitment.

### Gain of precursor gene expression

Intriguingly, 7% of *yy1f/f Mb1CRE* pro-B cells on OP9-DL4 feeders were identified as stem cells (Figure 5C and D), although the assignment did not appear to correlate with any specific cell cluster. We assayed for expression of two genes highly expressed in hematopoietic precursors and found elevated expression of CD34 and Itgax throughout the UMAP expression profile in the *yy1f/f Mb1CRE* pro-B cells grown on OP9-DL4 feeders (Figure S4A, panel 1) and very low expression within the three control samples (Figure S4A, panels 2-4). Elevated expression of these genes suggests a possible de-differentiation pathway of YY1-null pro-B cells into hematopoietic precursors.

In summary, our scRNA-seq results show that YY1 knock-out pro-B cells grown on OP9-DL4 feeders dramatically lose expression of transcripts representative of the B cell lineage profile, and simultaneously gain clusters of cells expressing numerous key genes indicative of various alternative hematopoietic lineages. These results indicate a loss of B cell lineage commitment, and development of an unusual plasticity of gene expression caused by loss of transcription factor YY1. Given sufficient time (3 weeks) and under defined conditions (OP9-DL4 feeders with IL7, SCF, and Flt3L), these cells adopt a phenotype highly consistent with T lineage cells.

### Some de-regulated genes show persistent expression after YY1-null pro-B cells develop into T-like cells

Our scRNA-seq data demonstrate dramatically increased expression of alternative hematopoietic lineage genes 2 weeks after YY1 knockout pro-B cells were incubated on OP9-DL4 feeders. While most of these alternative lineage genes are ultimately repressed as the cells transition to the T-like cell lineage (Figure 3), we reasoned that some alternative lineage genes might persist in expression, particularly if they are directly repressed by YY1. We evaluated expression of the top 25 genes (Table S1) from dendritic cells (DC4+, DC8+, DCpDC) and monocyes (6C+ 11-BI and 6C-11-BI) in our traditional RNA-seq data from wild-type thymic T cells, YY1-null pro-B cells differentiated into T-like cells, wild type pro-B cells, and YY1-null pro-B cells. This comparison showed elevated expression in YY1-null pro-B cells developed into T-like cells of dendritic genes Itgb7, Runx2, I830077J02Rik, Havcr2, Ccl5, and Ccr5, as well as monocytic genes Ifi204, Emilin2, and Tmem51 (Figure S4B and S5A panel 1). These genes were not highly expressed in control T cells or either of the pro-B cell populations (Figure S4B and S5A panels 2-4) but are highly activated during the transition from pro-B cells to T-like cells (Figure S4B and S5A panel 1).

Highest level expression was observed for the Itgb7 gene (Figure S4B and C; and Figure S5A panel 1). YY1 binds to this gene in pro-B cells at an intronic location (Figure S5B red oval) with properties of a potential DNA loop anchor as it also binds CTCF and Rad2. This DNA region also binds MED1, BRG1, IRF4, PU.1, and p300 implicating a transcriptional regulatory site (Figure S5B). Thus, YY1 may directly repress Itgb7 in pro-B cells, but as YY1-null pro-B cells begin to develop into T-like cells and the Itbg7 gene becomes activated (Figure S5A), the gene is ultimately not re-repressed in the developed T-like cells perhaps due to the absence of direct YY1 DNA binding (Figure S5B).

Monocyte genes Emililn2, Runx2, and Tmem51 also show elevated expression during transition from YY1-null pro-B cells into T-like cells (Figure S4B and S5A panel 1). These genes exhibit Ezh2 binding at their promoters in pro-B cells implicating a PcG mechanism of repression (Figure S5B red ovals). The ability of YY1 to initiate PcG repression has been demonstrated previously (Atchison et al. 2003) implicating a mechanism for the inability to re-repress these genes in the developed T-like cells. Overall, the elevated expression of alternative lineage genes revealed in our scRNA-seq data is matched in some cases by persistent expression of these genes after the cells develop from YY1-null pro-B cells into T-like cells. Regulation of some of these genes by YY1 may be direct (Itgb7 for instance) or indirect.

## Discussion

Our results show that knock-out of transcription factor YY1 in pro-B cells results in loss of B lineage commitment and gain of the plasticity to adopt the T-like cell lineage, and perhaps other hematopoietic cell lineages such as monocytes, dendrocytes, NK cells, and stem cells. We found development of YY1 knockout pro-B cells to the T-like cell lineage can occur *in vitro* using the OP9-DL4 system which promotes T lineage development, or *in vivo* by injection into sub-lethally irradiated *Rag1^-/-^* mice. The cells grown for three weeks *in vitro* possessed transcript profiles that closely align with the thymic T cell lineage, and nearly completely extinguish B lineage markers. This novel lineage plasticity is clearly the result of YY1 knockout as provision of YY1 expression by retroviral transduction eliminated this lineage plasticity. Interestingly, scRNA-seq experiments to assess this differentiation process mid-transition revealed cell clusters expressing a multiplicity of alternative hematopoietic lineage genes, whereas the B lineage transcript profile was nearly completely extinguished. Although B cell lineage plasticity has been observed following knockout of several lineage-specific transcription factors (Nutt et al. 1999; Horcher et al. 2001; Ivanova et al. 2002; Mikkola et al. 2002; Cobaleda et al. 2007; Nechanitzky et al. 2013; Somasundaram et al. 2015), YY1 is unique in being a ubiquitous transcription factor expressed in all cell types suggesting a potentially universal mechanism of lineage commitment.

B lineage development involves multiple steps of transcriptional priming to progress from multipotent progenitors to B lineage cells (Lin et al. 2010; Miyai et al. 2018). Initial priming and dynamic occupancy by EBF1 at B lymphoid enhancers, as well as functions by E2A and Foxo1 initiate the B lineage pathway (Lin et al. 2010; Lenaerts et al. 2022). Subsequently Pax5 is required for continued B lineage commitment (Nutt et al. 1999). A variety of studies have shown that both EBF1 and Pax5 are critical for B lineage development and commitment, and their knockout can result in loss of B lineage identity, and development into a variety of alternative hematopoietic lineages (Nutt et al. 1999; Horcher et al. 2001; Ivanova et al. 2002; Mikkola et al. 2002; Cobaleda et al. 2007; Nechanitzky et al. 2013; Somasundaram et al. 2015; Gruenbacher et al. 2023). Complete loss of Ebf1 and Pax5 expression in YY1-null cells on DL4 feeders indicates a critical role of YY1 in maintaining B cell identity, at least partially due to an important role in maintaining Ebf1 and Pax5 expression.

A growing body of evidence indicates that lineage initiation and commitment require not only activation of lineage specific genes, but also repression of genes expressed in alternative lineages. For instance, early in B lineage development, EBF1 binds to the *Gata3* promoter required for T lineage development, and in association with EZH2 represses *Gata3* expression (Banerjee et al. 2013). Many alternative lineage genes and enhancers are repressed by EBF1 and Pax5 to maintain B lineage fidelity (Mikkola et al. 2002; Cobaleda et al. 2007; Nechanitzky et al. 2013; Lenaerts et al. 2022). Similarly, TCF1, a pioneer factor for T lineage development, not only initiates expression of T lineage genes (Yui and Rothenberg 2014; Johnson et al. 2018), but also represses expression of GATA2 needed for mast cell development (Goldman et al. 2023). Additionally, deletion of the repressive transcriptional co-factor histone deacetylase gene HDAC7, leads to lymphopenia and B lineage promiscuity (Azagra et al. 2016) consistent with the importance of transcriptional repression for lineage development.

YY1 is well known for its ability to both activate and repress transcription and derives its name from these dual functions (Park and Atchison 1991; Seto et al. 1991; Shi et al. 1991). Its activation properties map to the N-terminal regions, whereas repression functions map to histone deacetylase binding regions, as well as to a small 25 amino acid segment that supports Polycomb-group repression (the REPO domain) (Bushmeyer et al. 1995; Austen et al. 1997; Galvin and Shi 1997; Thomas and Seto 1999; Satjin et al. 2001; Atchison et al. 2003; Srinivasan and Atchison 2004; Wilkinson et al. 2006; Basu et al. 2010). In addition, YY1 can self-associate providing a mechanism for it to bridge promoters and enhancers via long-distance DNA interactions which have been observed in B, T, neural, erythroid, and stem cell systems (Hwang et al. 2013; Medvedovic et al. 2013; Mehra et al. 2016; Beagan et al. 2017; Dong et al. 2022). Indeed, YY1’s function in long-distance DNA interactions is well documented in the B cell lineage where YY1 knockout leads to de-contraction of Ig loci and reduced V(D)J rearrangement of distal Ig genes (Liu et al. 2007). Similarly, deletion of YY1 in splenic B cells results in reduced Ig class switch recombination caused by a decrease in DNA loop formation between the IgH Eμ and 3’RR enhancers (Mehra et al. 2016). As enhancer-promoter loops often increase at key genes during lineage commitment (Hu et al. 2018; van Schoonhoven et al. 2020), it is possible that reduction of YY1-mediated promoter-enhancer DNA loops upon YY1 knockout reduces stable expression of genes required for B cell lineage commitment.

Our scRNA-seq data show that as YY1 knockout pro-B cells develop on OP9 DL4 feeders, they lose expression of B lineage transcripts, and concomitantly these cells exhibit an extraordinary increase in expression of genes characteristic of numerous alternative hematopoietic lineages. Rather than a gradual transition from a B cell to T cell phenotype as might be anticipated, we observed a much more diverse expression of a multitude of genes indicative of many alternative hematopoietic lineages. The large fraction of cells identified as monocytic or dendritic (85%) compared to the small fraction identified as T cells (3%) at this early time point suggests that it is more facile to convert YY1-null pro-B cells into these hematopoietic lineages compared to T cells. Incubation of YY1-null pro-B cells on OP9-DL4 feeders for a more extended time (3 weeks) results in down-regulation of most of these alternative lineage genes, and increased expression of the T lineage genes.

The appearance of clusters of cells representative of many alternative lineages argues that YY1 is required to repress key genes that regulate numerous alternative hematopoietic lineages. This is supported by continued expression of some alternative lineage genes after YY1 knockout pro-B cells fully develop into T-like cells after 3 weeks on OP9-DL4 feeders (Figure S4). The appearance of cells expressing stem cell transcripts (Figure S4A) also suggest that YY1 knockout pro-B cells may be capable of de-differentiating to a stem-like phenotype thus enabling development of numerous alternative hematopoietic lineages.

The multiplicity of cell types representing distinct hematopoietic lineages in our scRNA-seq experiments suggest that YY1 repression of alternative lineage transcripts may be a common mechanism in regulating lineage-specific genes. This, coupled with the positive impact of YY1 on expression of lineage-appropriate genes implies a mechanism of both YY1 transcriptional activation, as well as transcriptional repression in lineage commitment (Figure 8E). We propose that in wild-type pro-B cells, YY1 stably represses alternative lineage genes while simultaneously stably activating B lineage genes (Figure 8E, left panel). In YY1 knockout pro-B cells, B lineage gene expression is no longer stably enforced resulting in reduced expression of some B lineage genes, whereas alternative lineage genes are no longer stably repressed resulting in leaky expression. Incubation of these cells in a T cell inducing environment (OP9-DL4 cells for instance), thus leads to lost B lineage gene expression and gain of alternative linage gene expression (Figure 8E, right panel).

The above properties, coupled with its ubiquitous expression, suggest that YY1 is critical for commitment of multiple, and perhaps all, cell lineages representing a universal mechanism for lineage commitment. The appearance of cells with stem cell-like transcript profiles also suggests potential applications of transient YY1 knockdown for regenerative medicine approaches.

## Materials and methods

### Mice

C57BL/6 (CD45.1), Mb1CRE, C57BL/6 (CD45.2), and Rag1^−/−^ mice were purchased from the Jackson Laboratory. *yy1f/f* mice on a C57BL/6 background were a gift from Y. Shi (Oxford). *yy1f/f* mice were crossed with *Mb1-CRE* mice to generate *yy1^f/f^ Mb1-CRE* mice in which the endogenous *yy1* gene is conditionally knocked out at the pro-B cell stage by the action of Mb1-driven CRE recombinase. *yy1^f/f^ Mb1-CRE* mice and control mice between seven and nine weeks of age were used for all the experiments. All experiments involving animals were approved by the IACUC of the University of Pennsylvania and conform to the appropriate regulatory standards.

### Transplantation

For bone marrow transplantation, bone marrow cells were harvested from eight week-old *yy1f/f Mb1CRE* mice 4 days after intravenous injection of 250 μg/kg 5-fluorouracil (5-FU). Cells were cultured overnight in DMEM with 10% FBS, 1% antibiotic, 1% L-glutamine, IL-3 (10 ng/ml), IL-6 (5 ng/ml), and SCF (100 ng/ ml), then transduced with either empty retroviral vector (MigR1) or with vector expressing the YY1 cDNA (MigR1-YY1) by using polybrene (4 μg/ml) and the same cytokine cocktail. At least 1 million cells were injected intravenously into lethally irradiated (9 Gy) recipient CD45.2 mice. Antibiotic containing drinking water was provided for recipient mice for 2 weeks post transplantation. For pro-B cell transplantation, 400,000 to 500,000 sorted pro-B cells were injected intravenously into sub-lethally irradiated (450 rad) *Rag1^−/−^* recipient mice.

### Recombinant vectors and virus packaging

Flag-tagged YY1 cDNAs were cloned into the HpaI site of GFP-expressing MSCV-IRES-GFP vector (MigR1) by blunt end ligation. High titer retroviral supernatants were prepared following transfection of HEK293T cells. 293T cells were maintained in DMEM supplemented with 10% FBS, 1% Penicillin/Streptomycin, and 2 mM L-glutamine. Retroviral supernatant was then used for spin infection at 2500 rpm for 90 min in the presence polybrene (4 μg/ml). A second round of spin infection was performed 24 h following the first one.

### In vitro T cell differentiation

T cell differentiation assays were performed using OP9-DL4 stromal cells as described previously (Mohtashami et al. 2013). Briefly, OP9-DL4 stromal cells were maintained in MEMα nucleosides media containing 15% fetal bovine serum, 50μM β-mercaptoethanol, and antibiotic. Prior to initiation of co-culture, OP9-DL4 stromal cells were irradiated. Seven to nine-week-old *yy1^f/f^*or *yy1^f/f^ Mb1-CRE* mice were sacrificed, and bone marrow cells were collected. After red blood cell lysis, nucleated cells were blocked by Fc blocker, incubated with pro-B cell marker antibodies (B220, CD19, CD93, IgM, CD43, CD23) and sorted on a BD FACSAria™ Cell Sorter. Sorted pro-B cells (B220^low^/^+^CD19^+^CD93^+^IgM^−^CD43^+^CD23^-^) from *yy1^f/f^* or *yy1^f/f^ Mb1-CRE* mice were added to pre-established OP9-DL4 stromal layers and cultured in αMEM media supplemented with 10% FBS, β-mercaptoethanol, Penn-Strep, and 5 ng/ml IL-7, 5ng/ml Ft3L, and 10ng/ml SCF, respectively. After 7 days cells were re-plated on fresh irradiated OP9-DL4 cells. After approximately 3 weeks total in culture, non-adherent cells were harvested and analyzed by flow cytometry for the presence of Thy1.1 and CD25.

### Flow cytometric analysis and cell sorting

Single cell suspensions were stained and analyzed on an 18-color LSR fortessa flow cytometer (Becton Dickinson) equipped with four lasers for excitation of Blue, Red, Green, and Violet excited dyes. Antibodies were tagged with FITC, PE, PE-Cy5, PE-Cy7, APC or AF647, APC-Cy7, BV421, or BV605 versions of purified antibodies against CD3 (17A2), CD4 (RM4-5), CD8a (53-6.7), CD25 (PC61), CD44 (IM7), TCRβ (H57-597), Thy1.2 (30-H12), B220 (RA3-6B2), CD19 (6D5), CD43 (1B11), IgM (11/41), CD93 (AA4.1), CD23 (B3B4), c-kit (2B8), Sca-1 (D7), CD69 (H1.2F3), CD45.2 (104), and CD45.1(A20). All directly conjugated antibodies were purchased from eBiosciences (San Diego, CA), Bioegend (San Diego, CA), BD Pharmingen (San Diego, CA) or Invitrogen (Carlsbad, CA). DAPI and Zombie Aqua were used to stain dead cells. All files were analyzed with FlowJo software (Tree Star, Inc).

### PCR detection of YY1 deletion efficiency

Sorted CD25^+^ Thy1.2^+^ or CD4^+^ CD8^+^ cells (1 × 10^5^) were resuspended in 80 μL of 50 mM NaOH, heated for 5 min at 95°C, and vortexed to dissolve the cell pellets. NaOH was neutralized with 20 μl of 1 M Tris-HCl (pH 6.8). The resulting DNA solution was used for each PCR reaction to detect the deletion efficiency of the floxed *yy1* allele by using forward primer ACCTGGTCTATCGAAAGGAAGCAC and reverse primer CCAAAGTTCGAAACCTGCTTTCCT. PCR products were separated on 2% agarose gels and visualized by ethidium-bromide staining.

### Ig heavy chain V(D)J rearrangements analysis

CD25^+^ Thy1.2^+^ cells from *in vitro* differentiation cultures or CD4^+^ CD8^+^ cells from reconstituted thymocytes from Rag1^-/-^ mice were isolated using a BD FACSAria Cell Sorter. DNA was prepared by using 50 mM NaOH and 1 M Tris-HCl (pH 6.8) and semi-nested PCR was performed for detecting Ig heavy chain V(D)J rearrangements. Amplification processes were carried out in two rounds: Round 1 containing 5’primers VH 7183, or VHQ52, and JH4E primer (Key Resources Table). First round amplification was performed for 30 cycles (1min at 95°C, 1min at 60°C, 1.5 min at 72°C). For the second-round amplification, 1μl of the first-round product was used as a template and PCR consisted of 25 cycles (1 min at 95°C, 1 min at 63°C, 1.5 min at 72°C) with the same primers. Amplifications were performed with either primer pairs VH 7183 and the nested JH4A, or primer pairs VHQ52 and the nested JH4A (Key Resources Table).

### TCR beta chain V(D)J rearrangement analysis

Sorted CD25^+^, Thy1.2^+^ cells or CD4^+^ CD8^+^ cells were used to detect the TCRβ V(D)J rearrangements. DNA was prepared by using 50 mM NaOH and 1 M Tris-HCl (pH 6.8) as previously described. Nested PCR was performed in two rounds using the following primers: Vβ4 or Vβ8, and jβ2 and jβ2.7 nested (Key Resources Table). First round amplification was for 30 cycles (1 min at 95°C, 1min at 60°C, 1.5 min at 72°C) and second round amplification was for 20 cycles (1 min at 95°C, 1 min at 63°C, 1.5 min at 72°C).

### Bulk RNA-seq library preparation

The cDNA of sorted CD25^+^, Thy1.2^+^ cells either from *in vitro* differentiation cultures or from wild type mouse thymocytes were used to prepare sequencing libraries using the SMART-Seq® HT PLUS kit (R400748). Briefly, the quantification of total RNA was performed using a Qubit instrument, and the integrity of the RNA was confirmed using a TapeStation. Libraries were dual indexed and pooled according at equal molecular concentrations. Subsequently, 100-base pair reads were sequenced on an illumina HiSeq 2000 platform.

### Processing of bulk RNA-Sequencing data

Fastq files of raw sequence data for Control T DN cells (C) and T-like cells developed from YY1-null pro-B cells (T) samples were generated using Illumina bcl2fastq software. The quality of fastq raw reads was checked using FastQC (Andrews 2010). A total of 125,547,313 and 157,418,878 filtered reads were generated from YY1-null pro-B cells incubated for 3 weeks on OP9-DL4 feeders, and wild-type thymic T cells, respectively. The average filtered rate percentage was 85.49 and 83.41 and GC content was 43 and 44 in YY1-null pro-B cells incubated for 3 weeks on OP9-DL4 feeders, and wild-type thymic T cells, respectively.

To eliminate low-quality reads, paired end reads were filtered using the Trimmomatic 0.36 (Bolger et al. 2014) tool with average base quality < 20. After quality filtration, filtered reads were used for reference mapping using STAR(Dobin et al. 2013) (v.2.7.10a). The GRCm38 (mm10) mouse genome (http://useast.ensembl.org/Mus_musculus/Info/Index) was used for mapping.

Raw reads for control pro-B cell (B) and YY1-null pro-B cell (Y) samples were retrieved from National Center for Biotechnology Information-Sequence Read Archive (NCBI-SRA) with accession number PRJNA297235. The raw reads generated in the current study for Control T DN cells (C) and YY1-null T DN cells (T) were submitted to NCBI-SRA with accession number PRJNA903893. Unsupervised clustering analysis was performed using factextra (https://cran.r-project.org/web/packages/factoextra/index.html) and heatmaps were plotted using the gplots package in R.

### Differential gene expression analysis

The differential expression analysis of YY1-null T DN cells and Y1-null pro-B cells, with respect to their control samples, was done by estimating the total read count of assembled transcripts. The total read count was retrieved using StringTie (Pertea et al. 2015). Further these count values were used for differential expression analysis using the DESeq2 (Love et al. 2014) package in R Studio. The gene annotation of differentially expressed genes (DEGs) was carried out with GRCm38 (mm10) mouse genome (http://useast.ensembl.org/Mus_musculus/Info/Index). The threshold for significant differential gene expression was set as log2 fold change (FC) ≥ 1 and ≤ −1 with ≤ 0.05 p-value for up-regulated and down-regulated genes.

### Gene ontology (GO) and Kyoto encyclopedia of genes and genomes (KEGG) pathway enrichment analysis

To study the putative functions of the identified differentially expressed genes, gene ontology (GO) analysis was done that classified genes into cellular component (CC), molecular function (MF), and biological process (BP) categories. GO enrichment analysis was performed using org.Mm.eg.db keytype and the enrichplot package in RStudio with < 0.05 p-value cutoff. To understand the enriched pathways of significant differentially expressed genes, KEGG analysis was performed. KEGG pathways analysis was employed using the clusterProfiler package (Yu et al. 2012) with the number of permutations (nPerm) set to 10000, org.Mm.eg.db and pAjustMethod set to “none”. P-value ≤ 0.05 was set for significant KEGG pathways.

### Sc-RNA-seq library preparation

Biological replicates for scRNA-seq from mouse bone marrow cells were isolated from *yy1^f/f^* and *yy1^f/f^ Mb1CRE* mice and FACS-sorted for pro-B cells as described. Cells were either subjected to scRNA-seq directly or were cultured on OP9-DL4 feeders as described above for 12-14 days when the *yy1^f/f^ Mb1CRE* cells had become 3 to 5% Thy1.2^+^ CD25^+^. Next-generation sequencing libraries were prepared using the 10x Genomics Chromium Single Cell 3’ Reagent kit v3 according to the manufacturer’s instructions. Libraries were uniquely indexed using the dual index kit, pooled, and sequenced on an Illumina NovaSeq 6000 sequencer in a paired-end, dual indexing run. Sequencing for each library targeted 20,000 mean reads per cell.

### scRNA-seq data analysis

Raw data was then processed using the Cell Ranger pipeline (10x Genomics, v.6.1.2) for demultiplexing with the mkfastq command to generate Fastq files and for alignment of sequencing reads to the mm10 reference genome and creation of feature-barcode matrices. Secondary data analyses were performed in R (v.4.2.2) with Seurat (v4.3.0). The unique molecular identified (UMI) count tables were initially loaded into R (v4.2.2) using the Read10X_h5 function, and then Seurat objects were created for each sample. The samples were merged, and during the normalization process in Seurat, SCTransform was used to regress out the effects of library size. In Seurat, the cells were clustered using a clustering algorithm based on shared nearest neighbor modularity optimization and dimensional reduction was performed with first 30 PCs. Cells were annotated with Immgen data using the SingleR (v1.8.1) package and marker gene expressions were shown with FeaturePlot and stacked violin plot using Seurat (v4.3.0), cowplot (v1.1.1) and ggplot2 (v3.4.0) packages in RStudio. The count of each cell type was retrieved using the tensorflow (2.14.0) and celldex (v1.4.0) packages.

### Key Resources Table

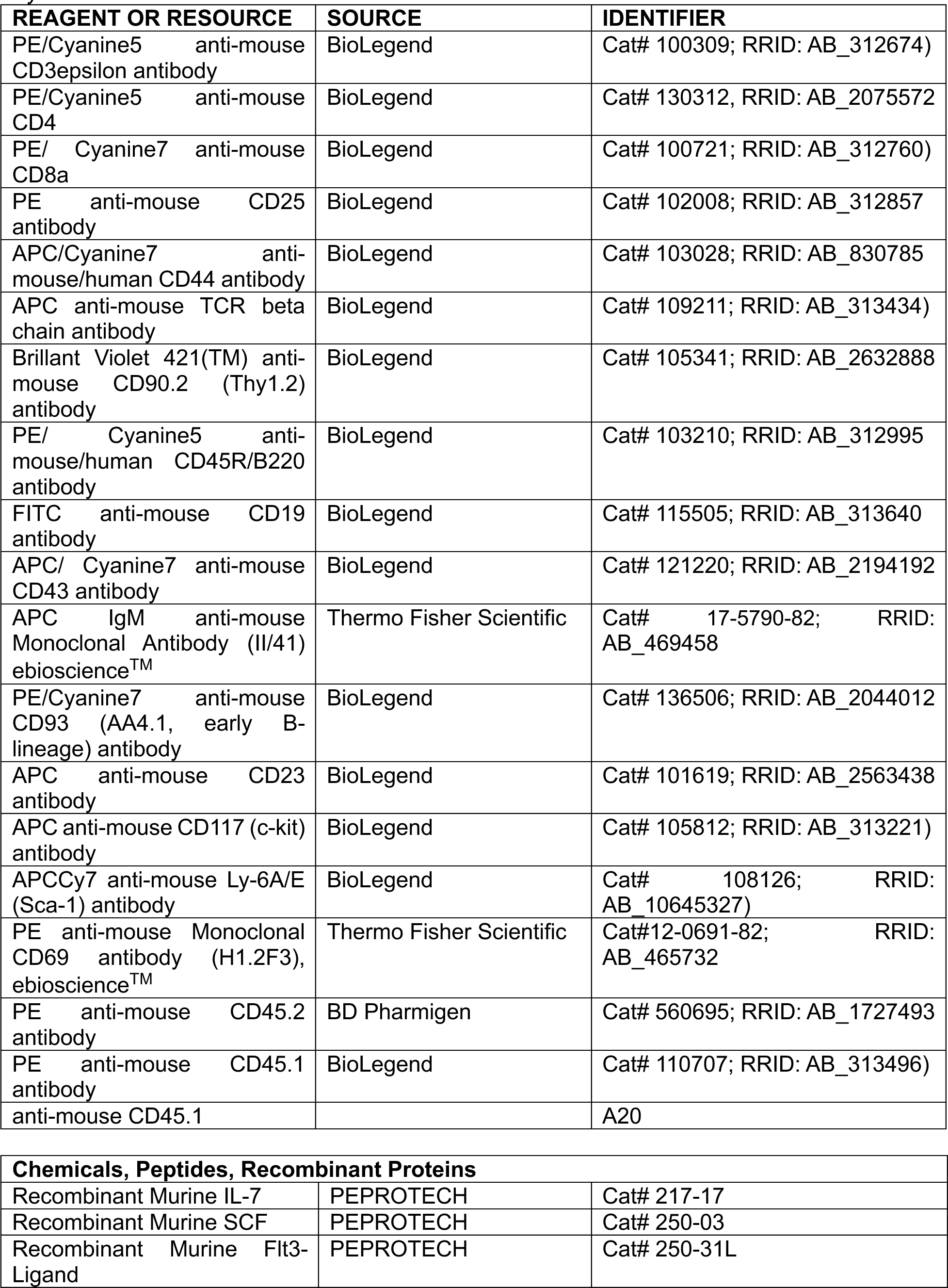
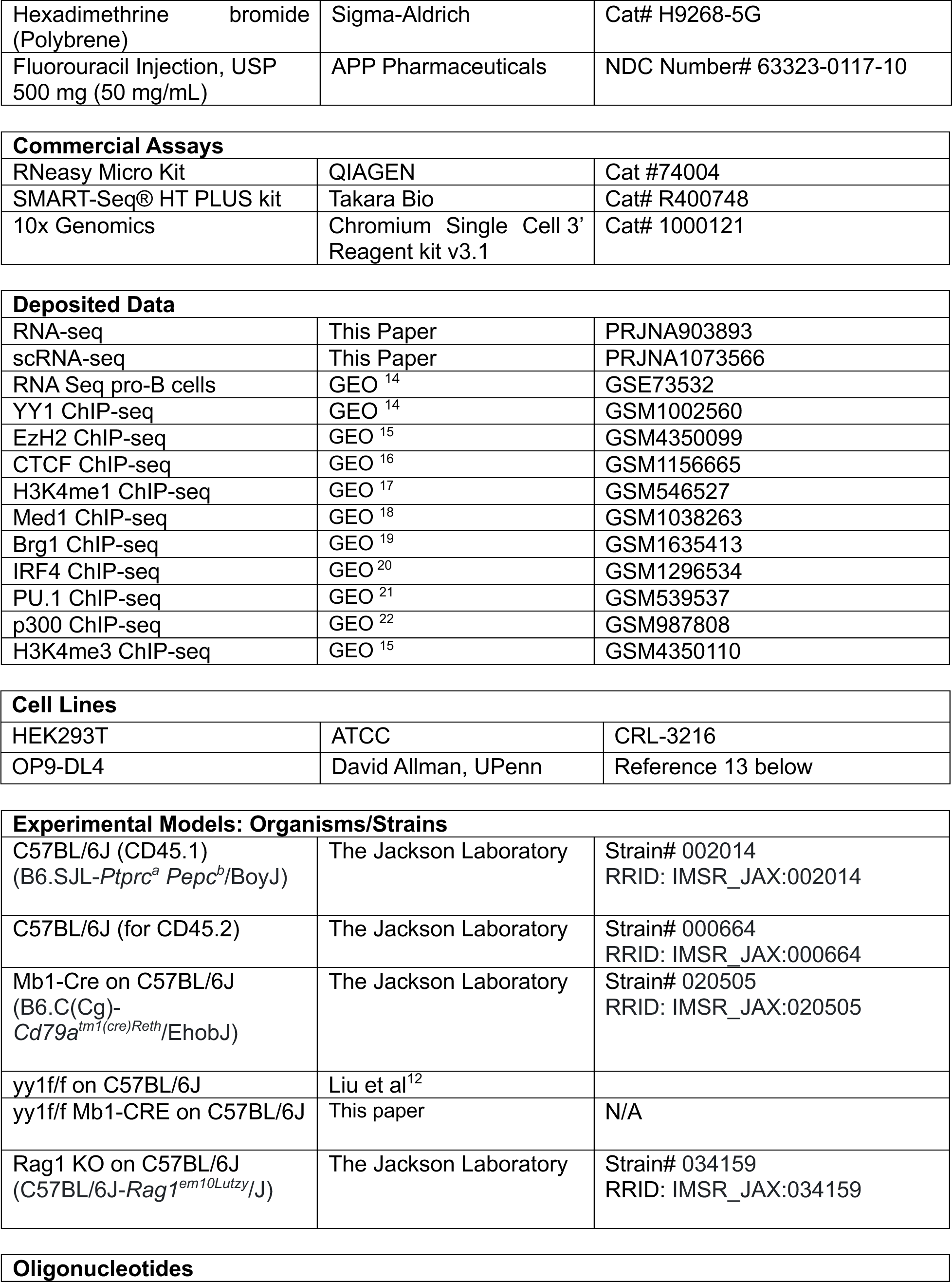
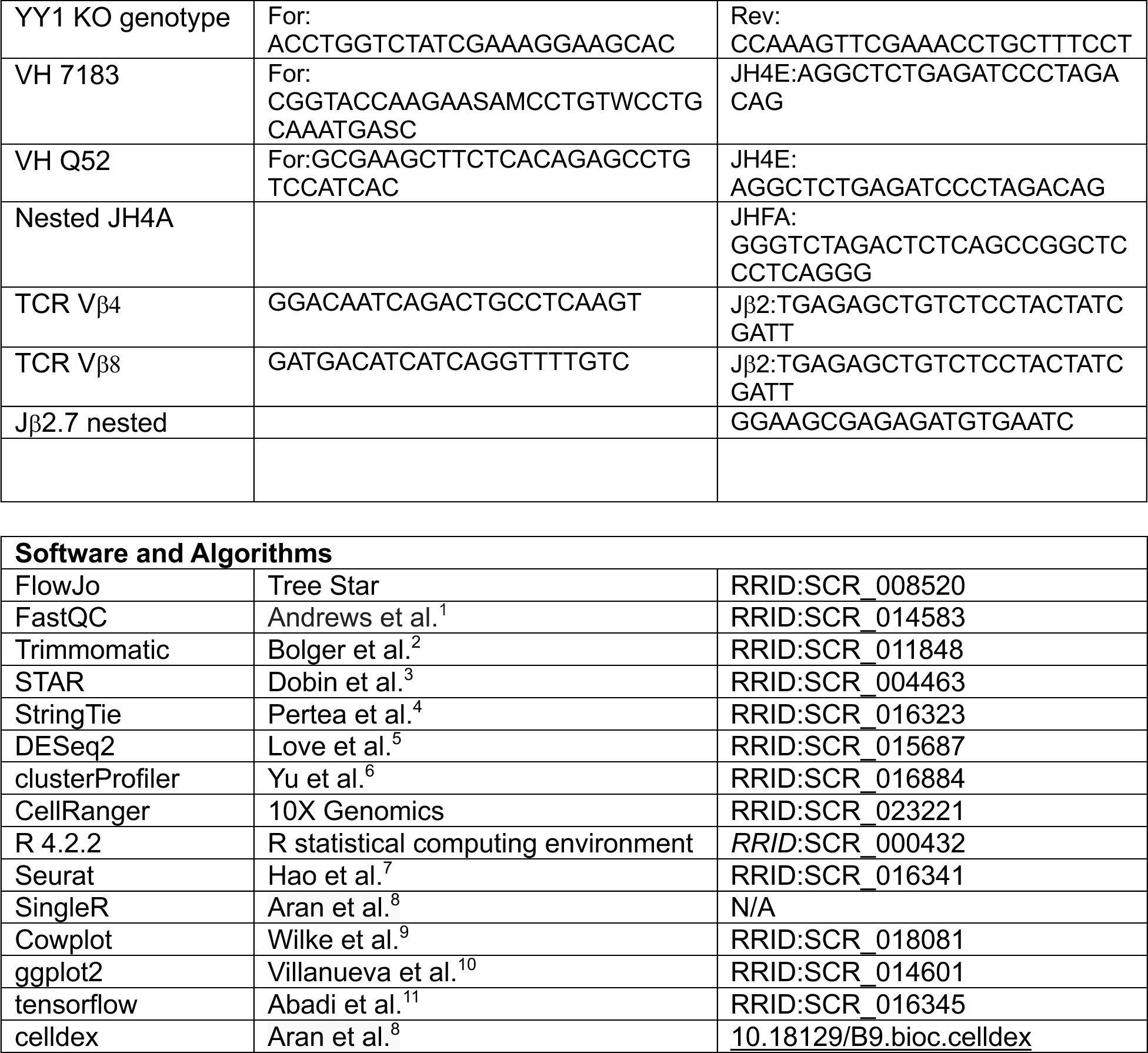

## Author Contributions

MLA and DA conceived the study, MLA supervised the study, SB, SS, SN, SH, AB, NB, and JR designed and performed experiments, SB, SS, SH, SN, NB, AB, JR, DD evaluated data, MLA, SS, SN, NB, DD, prepared figures, MLA, SB, SS, SN, NB, and SH wrote the manuscript.

## Acknowledgements

This work was supported by NIH grants R01AI162879 and R01AI155540 to MLA and grants R01AI139123, R01AI175185, and an Aspire award from the Mark Foundation for Cancer Research to DA. We are thankful for the Penn Cytomics and Cell Sorting Shared Resource, the School of Veterinary Medicine Cytometry Core, the Center for Host-Microbial Interactions sequencing facility, the Penn Next Generation Sequencing Core, the Children’s Hospital of Philadelphia Single Cell Technology Core, and the Penn Genomic and Sequencing Core. Figures were prepared using Biorender.com.

---

**Table S1.**
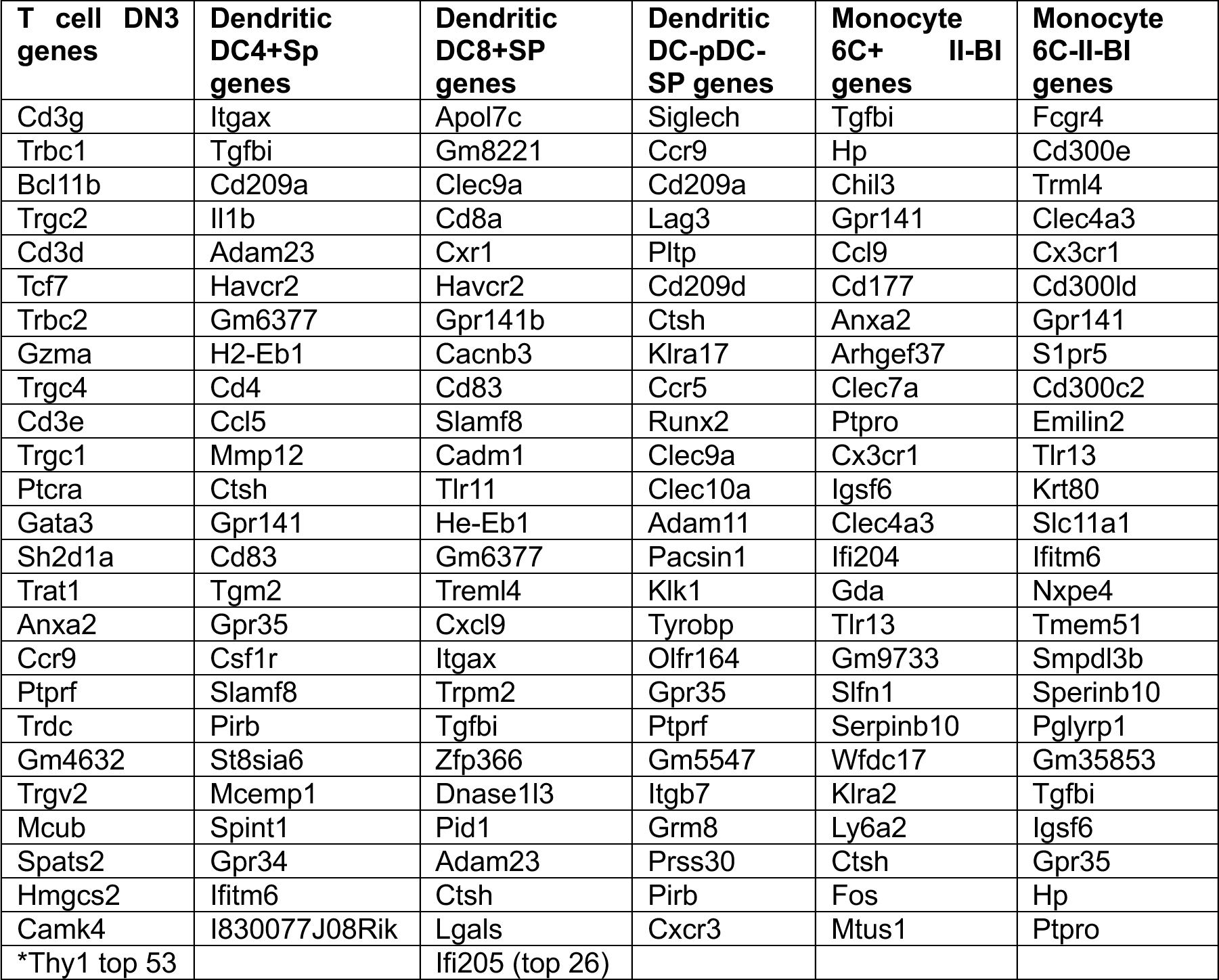
Top 25 Genes in Various Lineages Compared to pro-B FrB/C Cells. Immgen (https://www.immgen.org/) RNA-seq data from various purified hematopoietic cell types shown in the table were each run through the MyGeneset program against pro-B fraction B/C RNA-seq data. The top 26 to 26 genes enriched in each sample compared to pro-B fraction B/C cells is shown.

